# Single-cell RNA sequencing of human, macaque, and mouse testes uncovers conserved and divergent features of mammalian spermatogenesis

**DOI:** 10.1101/2020.03.17.994509

**Authors:** Adrienne Niederriter Shami, Xianing Zheng, Sarah K. Munyoki, Qianyi Ma, Gabriel L. Manske, Christopher D. Green, Meena Sukhwani, Kyle E. Orwig, Jun Z. Li, Saher Sue Hammoud

## Abstract

Spermatogenesis is a highly regulated process that produces sperm to transmit genetic information to the next generation. Although extensively studied in mice, our current understanding of primate spermatogenesis is limited to populations defined by state-specific markers defined from rodent data. As between-species differences have been reported in the process duration and cellular differentiation hierarchy, it remains unclear how molecular markers and cell states are conserved or have diverged from mice to man. To address this challenge, we employ single-cell RNA-sequencing to identify transcriptional signatures of major germ and somatic cell-types of the testes in human, macaque and mice. This approach reveals differences in expression throughout spermatogenesis, including the stem/progenitor pool of spermatogonia, classical markers of differentiation, potential regulators of meiosis, the kinetics of RNA turnover during spermatid differentiation, and germ cell-soma communication. These datasets provide a rich foundation for future targeted mechanistic studies of primate germ cell development and *in vitro* gametogenesis.

## Introduction

Sperm are highly specialized terminally differentiated cells that carry the genetic information from father to offspring, thus providing a continuous link between the past and future of a species. In all male mammals, the foundational unit of fertility is the spermatogonial stem cell, which balances self-renewal with differentiation to sustain continuous sperm production throughout a male’s adult life. In all mammals, isolated spermatogonial stem cells divide to form networks of connected cells which progress through a series of mitotic and meiotic divisions followed by post-meiotic morphological changes of spermiogenesis to produce mature sperm. This differentiation process is reliant on intrinsic (germ-cell mediated) and extrinsic (soma mediated) signals. Despite these global similarities, it is difficult to translate knowledge generated in mice to higher primates, especially as it relates to spermatogonial development, because the classical vocabulary that is used to describe those cells in rodents [A_single_ (A_s_), A_paired_ (A_pr_), A_aligned_ (A_al_), A1-4, Int, B]^1–3^ is different than monkeys (A_dark_, A_pale_, B1-4) and humans (A_dark_, A_pale_, B)^4–7^ For example, based on the existing terminology, it may not be obvious that the undifferentiated to differentiated transition that is marked by expression of cKIT corresponds to the Aal to A1 transition in mice while it corresponds to the A_pale_ to B transition in monkeys and humans^8^.Therefore, identification and direct comparison of equivalent cell types and states is currently lacking in the field, and as a result, there has been limited success in applying knowledge from mice to inform the study of spermatogenesis and fertility in higher primates.

Although the basic cellular events underlying spermatogenesis and the ultimate goal of gamete production remain the same, there are numerous interspecies differences, including the histological organization of the testis, duration of the seminiferous epithelium cycle, dynamics of stem cell renewal and differentiation, and the quantity and morphology of sperm produced (reviewed in^9–11^). Therefore, to molecularly dissect the spermatogenesis process, translate knowledge gained between species, and to better make use of the diverse tools available in genetic and physiologic models like mice and macaques we needed a comprehensive and unbiased analysis of the gametogenesis process. Using the power of single cell transcriptomics, we sought to better discern molecularly analogous populations in order to identify shared and divergent properties of spermatogenesis between species, within both the germline and the soma.

Specifically, we analyzed ~14K adult human and ~21K macaque testicular cells, and combined these new datasets with our previously published mouse single cell data^12^. This allowed us to directly define homologous cell types across species using hundreds to thousands of genes, and uncovered both known and underrepresented cell types, including transient cell states too rare to be detected with low-throughput approaches. As a result, we produced a high-resolution three-species atlas with aligned analogous somatic and germ cell types/states. First, we describe six molecularly defined consensus spermatogonia states (SPG states) that are conserved in mice, monkeys and humans. While the early terminology used to describe undifferentiated spermatogonia in rodents (A_s_, A_pr_, A_al_) are different than monkeys or humans (A_dark_, A_pale_), the SPG cluster identities now provide a harmonizing vocabulary to facilitate crossspecies comparisons. Furthermore, we demonstrate that TSPAN33, a cell surface marker of SPG1, can be used to isolate and enrich transplantable human spermatogonia. However, a substantial number of transplantable human spermatogonia were also recovered in the TSPAN33 negative fraction, indicating that cells with colonizing potential in the transplant assay are heterogeneous, at least with respect to TSPAN33 expression. Additionally, we generate the first mammalian spermatogenesis pseudotime map, and describe several conserved and diverged molecular pathways operating within or across species. Finally, we describe between-species differences in somatic cell type relatedness/plasticity, functional roles, and their potential communications with the germline. Overall, this study provides the first single-cell comparative analysis of the spermatogenesis program between primates and rodents. Such a new resource is expected to improve our knowledge base for future studies of germ cell development in primates, and ultimately improve our understanding of the intrinsic and extrinsic evolutionary changes of the gametogenesis program. Knowledge gained from these data will inform fertility restoration efforts, including SSC culture and *in vitro* gametogenesis.

## Results

### Single-cell sequencing identifies major germ and somatic cell types of adult human and macaque testes

Using the Drop-seq platform, we generated single cell transcriptome data from ~14K and ~22K adult human and rhesus macaque testes, respectively. Using these datasets, our overall strategy was to first identify major cell types and states in the adult human and rhesus macaque testes separately, then combine them, along with our previous reported mouse data^12^, to define evolutionarily conserved or diverged programs within germ cells and the somatic cells of the testis (**Figure 1A**). The doublet rates for the human and macaque datasets is estimated to be <2%, therefore, allowing reliable analysis of individual cells (**Figure S1A**). After QC filtering (see Methods), 13,837 and 21,574 cells were retained from human and macaque, respectively, with an average of 3,210 unique molecular identifiers (UMIs) per cell, and 1,304 genes detected per cell. This sequencing depth and the number of detected genes was sufficient to define major cell types^13,14^. Systematic comparisons of technical batches (such as Human 1.1-1.5, **Figure S1B-C**) and biological replicates (4 humans and 5 macaques, **Figure S1D-E**) confirmed high batch-to-batch or individual-to-individual concordance, despite the variable proportions of cell types in different samples, likely due to differences in processing or storage of the tissue fragments analyzed.

**Figure 1.**
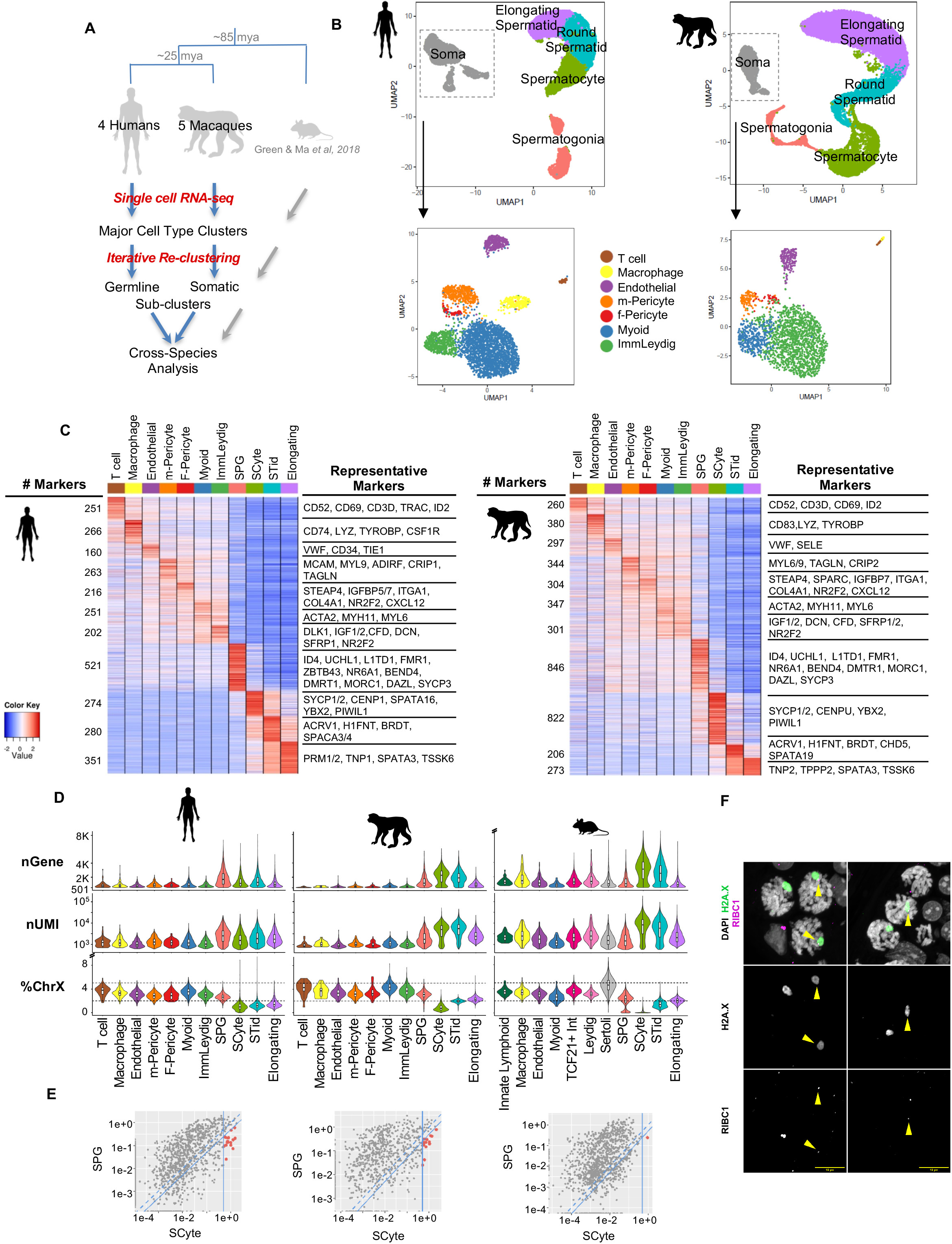

By unsupervised clustering of human and macaque cells separately, we defined 11 major cell types in both species. Using the expression pattern of previously known cell type-specific markers, we identified somatic cells and the four major germ cell types: spermatogonia (SPG), meiotic spermatocytes (SCytes), post-meiotic haploid round spermatids (STids) and elongating spermatids (Elong) (**Figure 1B-C, Supplemental Table 1**). Focused re-clustering of somatic cells defined seven major somatic cell types: macrophages, T cells, endothelial cells, peritubular myoid cells, two pericyte subpopulations (with either smooth muscle or fibroblast properties), and a Leydig cell precursor population (**Figure 1B insert, 1C, Supplemental Table 1**). Altogether, our atlas captures the majority of cell types in both species and allows us to directly compare testis cell types/states across human, macaque, and mouse to define recurrent or divergent molecular programs spanning 85 million years of mammalian evolution.

### Between-species differences of cellular attributes defined by transcriptomic properties

To better understand the global molecular and programmatic differences between primate and rodent germ and somatic cells, we compared several cellular attributes extracted from our data using transcriptome-defined indices (**Figure 1D**).

Previous comparisons of bulk RNAseq data across tissues have shown that testis, and particularly germ cells, express a relatively high number of detected genes^15,16^. To what extent this diverse gene expression reflects truly higher levels of unique transcripts per cell or rather a greater heterogeneity of distinct cell types in primate testis was not clear. Our scRNAseq data revealed that individual germ cells tend to express a larger number of genes (“nGene” in **Figure 1D**) than somatic cells in both macaque and human testis, consistent with our early findings in mice^12^. Interestingly, the largest diversity of expressed genes is variable across species: highest in spermatocytes and spermatids of macaques and mice, and in spermatogonia for humans (**Figure 1D**). Therefore, germ cell pervasive transcription occurs in different spermatogenic stages across species, although the functional significance of this germline attribute remains to be determined.

As germ cells tend to have higher number of total transcripts (“nUMI” in **Figure 1D**) sequenced per cell, this elevated “library size factor” could partially account for the higher number of genes detected. To discern the distribution pattern of transcripts within individual cells, we calculated a Gini Index for each cell and compared across cell types in all three species. This analysis quantified the level of transcriptional inequality among genes, determining whether a cell expresses many distinct genes at comparable levels or concentrates most transcripts on a small number of genes. At a given nUMI, a larger Gini index indicates a more biased distribution. Similar to what we previously observed in mice, the somatic cells and the spermatogonia population in human and macaque have the lowest Gini index, which increases progressively in the remaining germ cell populations, reaching the highest level (i.e., the most uneven distribution) in the elongated spermatids (**Figure S1F-G**). This trend indicates that germ cells along the differentiation trajectory devote an increasingly higher fraction of their transcriptome to a narrower set of genes, which likely reflects the focus on increasingly specific biological functions^17,18^.

During meiosis, sex chromosomes become transcriptionally silenced via meiotic sex chromosome inactivation (MSCI)^19^. In all three species, X-linked genes account for 2-5% of the cell’s total transcripts in the somatic cells and spermatogonia, yet this fraction is markedly lower in spermatocytes, and partially recovers in the round and elongated spermatids, although to varying degrees in different species (“%X” in **Figure 1D**). Furthermore, the suppression of sex chromosome gene expression in spermatocytes appears to be less complete in primates than in mice ((“%X” in **Figure 1D, Figure S1H**), and is truly limited to X-chromosome genes since non-X chromosome average gene expression in spermatocytes and the average X-chromosome gene expression in germ cell states proceeding MSCI are comparable **(Figure S1H)**. To generate a comprehensive list of genes reliably “expressed” in spermatocytes that escape MSCI in either species, we used the following two selection criteria: (1) absolute expression level cutoff: nUMI>0.5 in spermatocytes, and (2) two fold higher expression in spermatocytes than in spermatogonia to ensure that the escapee genes are being actively transcribed, rather than being protected from degradation (**Figure 1E**). Using this selection criteria, we identified 16 human, 14 macaque genes that escape MSCI, whereas the same threshold in mice identified a single gene Tsga8. These predicted escapees in each species do not enrich in the pseudo autosomal regions, rather they are scattered along both arms of the X-chromosome and lack an expressed Y-chromosome homologue (**Supplemental Table 1, see Discussion**). Of the genes that escape MSCI in primate spermatocytes, 7 are shared between the two species (**Supplemental Table 1, see Discussion**). To experimentally test whether these genes can truly escape MSCI, we designed Intron-seq fluorescent in situ hybridization probes for RIBC1 (a predicted escapee in humans and macaque), and show that RIBC1 is indeed transcribed in the XY body of pachytene spermatocytes of macaques (Figure 1F). This incomplete silencing of sex chromosomes observed in our human/macaque single cell data and now in macaque tissue cross-section is consistent with earlier cytological data showing that XY body in human pachytene spermatocytes maintain low levels of Cot1 and tritiated uridine staining in humans, whereas, Cot1 staining was undetectable in mice^20,21^. Therefore, our single cell data extend these cytological observations and identify candidate genes escaping MSCI.

Taken together, our global molecular analysis of germ cells and somatic highlights important differences in organization and/or regulation of the spermatogenic program among mammals.

### Iterative clustering of germ cells independently in each species reveals analogous patterns of discrete and continuous developmental trajectories

Previously, we re-clustered of 20K mouse germ cells and identified a single discrete developmental transition after spermatogonia (SPG) and a continuous developmental trajectory from spermatocytes to elongated spermatids^12^. Here, we extract 6,015 germ cells in human and 12,799 germ cells in macaque for focused re-clustering separately in each species, and similarly observed the clear partition between SPG and non-SPG germ cells for both species using either principal component analysis (PCA) (**Figure S1I-J**) or rank correlation among the seven clusters (**Figure S1K-L**). A provisional clustering analysis divides the germ cells into seven clusters for each species (hGC1-7 and mGC1-7) (**Figure S1H, I**). Using the expression of known cell state markers, we annotate hGC1-2, mGC1-2 as spermatogonia, hGC3 and mGC3-4 as meiotic spermatocytes, hGC4-5, mGC5 as post-meiotic round, and hGC6-7, mGC6-7 as elongated spermatids (**Supplemental Table 2**). While this seven-cluster solution will be immediately updated by finer partitions when we zoom in to the SPGs and non-SPG germ cells separately (see below), this mid-level analysis of germ cells provides independent catalogs of cell types/states and comprehensive lists of molecular markers for human and macaque that have not been restricted to 1:1:1 orthologous genes.

### Provisional clustering of SPGs from each species identifies four spermatogonial states

To get a cursory view of spermatogonia subtypes we further divided human SPGs (cells in hGC1-2) into finer groups by re-clustering, which uncovers four transcriptionally distinct states, hSPG1-4 (**Figure S1I, inset, Supplemental Table 2**). These four molecular states are highly correlated to the transcriptional states 0-4 (r = 0.83 - 0.93, **Figure S2A**) recently described by Guo *et al* and others^22–24^ Similarly, re-clustering of macaque spermatogonia (mGC1-2) finds four states referred to as mSPG1-4 (**Figure S1J, inset, Supplemental Table 2**). A similar analysis of our mouse spermatogonia also identifies four molecular states. The repeated identification of four states in each individual species raises the immediate question as to whether the four provisionally divided spermatogonial states, identified independently in each of the three species, map to each other in a one-to-one fashion. In the past, a common practice was to align cellular states across species based on a small selection of markers. However, here we have the opportunity to perform transcriptome-wide alignment to identify equivalent cellular states.

### Joint analysis of human, macaque, and mouse spermatogonia uncovers six analogous molecularly states

To directly identify comparable molecular states across species we used 1-1-1 homologous genes to merge the 1688 human SPGs, 747 macaque SPGs, and 2174 mouse SPGs. Compared to single-species analyses, this three-species merging significantly increases the total number of SPG cells, allowing more sensitive detection of rarer cell states that may have been missed in individual species. The re-clustering of ~4500 spermatogonia cells identifies six consensus states, ordered as SPG1-6 (**Figure 2A, S2B**). Expression patterns of previously established conserved marker genes suggest that SPG1/2 are undifferentiated spermatogonia, while SPG3-5 represent progressive stages of differentiation, up to SPG6 which consists of cells preparing for meiotic entry (**Supplemental Table 3**).

**Figure 2.**
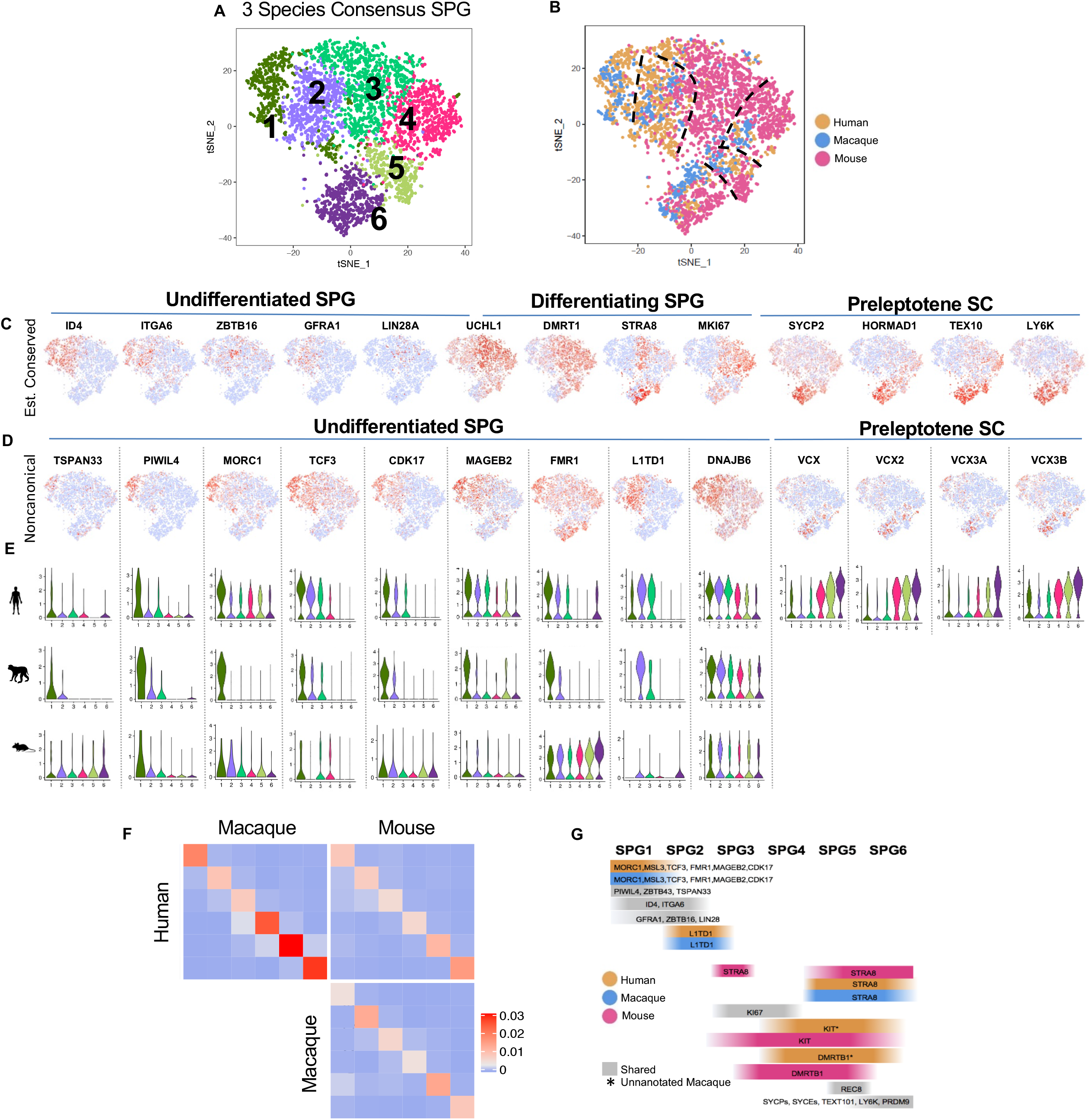

A closer examination of the composition of SPG1-6 states shows that the three species are not distributed evenly across the six consensus states (**Figure 2B, S2C, Figure S2D upper panels**). By tallying the cells’ initial assignment to the four SPG states defined in each species and to the six consensus states, we see that while the cells largely maintain their developmental ordering, there are notable between-species differences of state occupancy (**Figure S2D lower panels**). Notably, we find that the majority of cells in the undifferentiated SPG1 and SPG2 states are from macaque and human while fewer mouse cells occupy those states. In contrast, the majority of cells in differentiating SPG3 and SPG4 states are from mouse and fewer macaque and human cells occupy those states. All three species are represented in SPG5 and SPG6. The between-species shifts in state occupancy reflect differences in the relative size of the stem/progenitor pool and differing number of transit amplifying divisions (**Figure S2C-E**), as previously observed by histology (reviewed in^9^).

### Consensus SPG1-2

Although cells in SPG1 and SPG2 are similar to each other (**Figure S2B**) and at first glance express many of the well-established canonical markers of undifferentiated spermatogonia, (e.g., GFRA1, ZBTB16 (PLZF), ITGA6, ID4, UCHL1, LIN28, DPPA4 (macaque), FGFR3 and UTF1 (human) (**Fig 2D, S2F, Table S3**)^9,25–28^ the cells within each cluster enrich for a unique subset of canonical and noncanonical markers that can be used to clearly distinguish these two clusters (**Figure 2C,D,E**). Specifically, cells in SPG1 are largely contributed by human and macaque (**Figure 2B, S2C**), and are enriched for MORC1, MSL3, ZBTB43, TSPAN33, TCF3, FMR1, MAGEB2 and CDK17 (**Figure 2E**(TSPAN33, PIWIL4, MORC1;see **Table S3** for complete list of shared and species-specific markers). Interestingly, PIWIL4 is a significant SPG1 state-specific marker for all three species (**Figure 2E; Table S3**). However, mouse contributes very few cells to SPG1 (~40 cells; **Figure S2C**), accounting for ~2% of all mouse spermatogonia, which are too rare to have clustered independently in our earlier work. This observation is consistent with the understanding that the stem/progenitor pool of spermatogonia in mice is much smaller than in primates (reviewed in^9^.) In SPG2 cells, the noncanonical markers enriched in SPG1 (described above) are diminished, whereas, certain canonical undifferentiated spermatogonia markers (i.e. ZBTB16 (mouse and human only), GFRA1 (all three species)) increased in expression, as did several noncanonical markers (DUSP6, TCF4, L1TD1 (macaque and human only))(**Figure 2D,E**(L1TD1 shown); **Table S3**). Given these distinct markers, we conclude that consensus clusters SPG1&2 correspond to two distinct undifferentiated spermatogonia populations.

To validate newly-identified noncanonical primate markers expressed by SPG1 and SPG2 undifferentiated spermatogonia (TCF3, MAGEB2, CDK17, MORC1, DNAJB6, PIWIL4 and FMR1), we co-stained with UCHL1, a broad undifferentiated spermatogonia marker (**Figure 3A-D, S3A-E, upper panels**), or CKIT, a differentiating spermatogonia marker (**Figure 3A-D, S3A-E, lower panels)** in human testes tissue. TCF3 and PIWIL4 have been described in previous papers^22,23,28^, while MAGEB2, CDK17, MORC1, DNAJB6, and FMR1 are new markers described in this study. Consistent with our scRNAseq data predictions, our protein markers exhibit overlap with UCHL1+ spermatogonia cells lining the basement membrane of the human testis (**Figure 3A-D, upper panels**), but less overlap with cKIT+ differentiating spermatogonia (**Fig 3A-D, lower panels**); confirming that SPG1 and SPG2 are undifferentiated spermatogonial cell populations.

**Figure 3.**
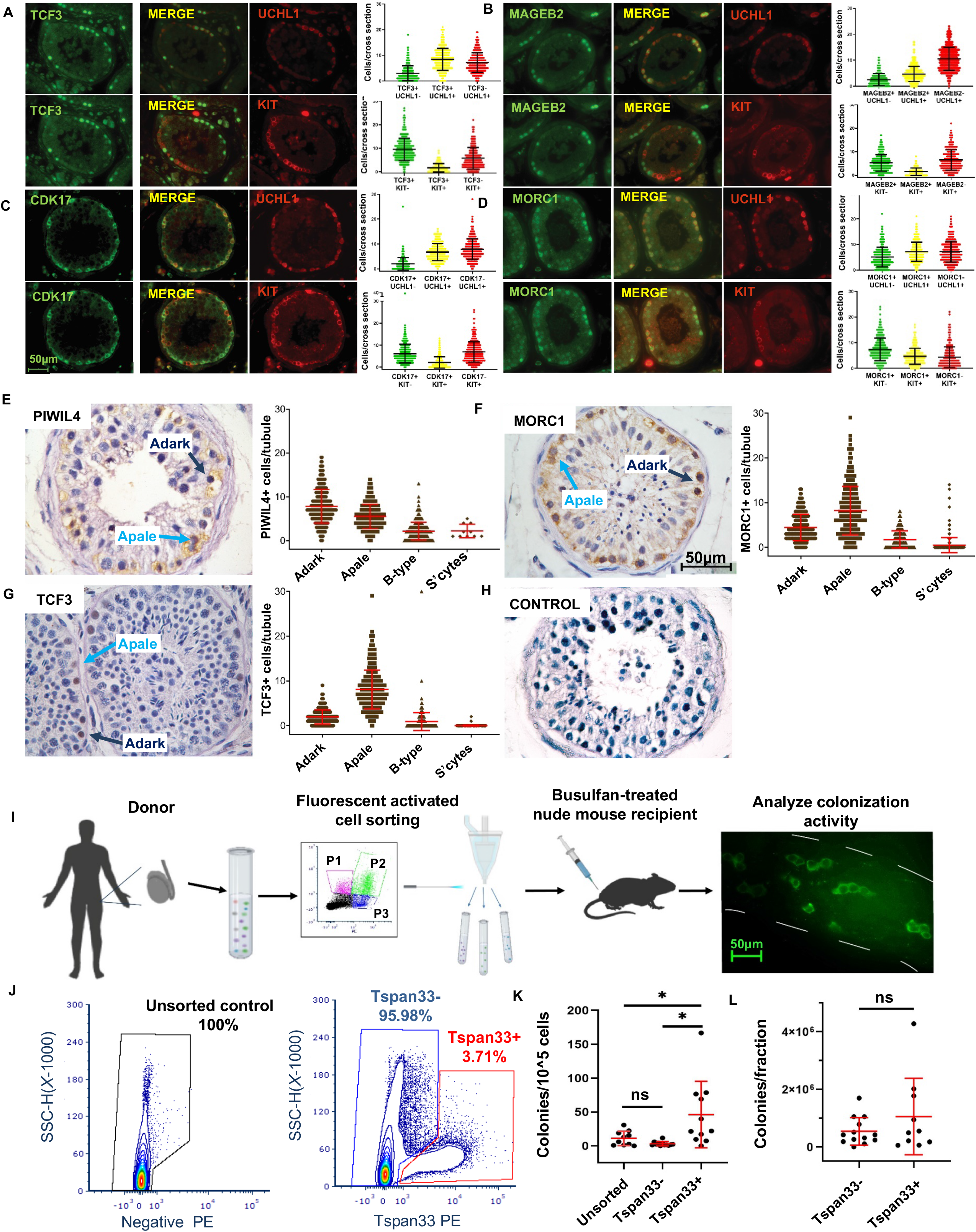

Next, we explored whether SPG1 and SPG2 represent cells in different phases of the mitotic cycle. We used ~500 genes known to vary with cell cycle^29^ to determine each cell’s likely phase (**Figure S3F-G**) and found that the clusters similarly enriched for cells in M/G1 and G1/S (**Figure S3H**). Thus, cell cycle state does not account for the difference between SPG1 and 2, but rather these molecular clusters define two transcriptionally distinct states of undifferentiated spermatogonia.

### Consensus SPG3-5

A combination of markers is often used to identify differentiating spermatogonia populations across species^9,28^. States SPG3-SPG5 are more similar to each other than to SPG1-2 or SPG6 (**Figure S2B**). Human and macaque testes contain cells that express many similar molecular markers to mouse SPG3 such as *DMRT1*, *KIT*, *STRA8, SOHLH1* and *SOHLH2 (*possibly equivalent to A_aligned_ - A1 spermatogonia in mice, A_pale_ - B1 macaque; A_pale_ - B human), but many of these markers are not restricted to specific developmental state – rather are broadly expressed in multiple differentiating SPG clusters and are often temporally uncoupled across species (**Figure S2F**, summarized in **Figure 2G**). For example, KIT positive cells are sporadic in SPG1-3 and then increase in frequency in human SPG4 (Macaque KIT not annotated), while STRA8 is absent in SPG3, but sharply upregulated in human and macaque SPG5-6. MKI67 is upregulated in SPG3 of mice and macaques, but not until SPG4 in humans (**Figure S2F**; **Table S3**). Cells in macaque and human SPG4 expressed markers such as MKI67, PCNA, numerous cyclins, DMRT1, DMRTB1 and KIT (unannotated in macaque), indicating that this population is actively proliferating, and appears to be molecularly analogous to cells transitioning from Type A1-4/ A_Intermediate_ to B spermatogonia in mouse (see **Table S3, Figure S2F**)^30^. This transition has a distinct cell cycle program depicted in **Figure S3F and G**, which appears to have either extended S/G2-phase or more transient G1 phase, possibly reducing the likelihood of premature meiotic entry of germ cells. The macaque and human SPG5 population continue to express DMRT1 and DMRTB1 (unannotated in monkey) while turning down KIT and turning on STRA8 as well as the meiotic gene REC8 (See **Table S3, Figure S2F**). Together, these data suggest that consensus SPG4 and SPG5 may represent two transcriptionally distinct Type B spermatogonia populations – referred to here as early (SPG4) and late (SPG5) Type B – in all three species. Classical descriptions of spermatogenic lineage development describes six populations of differentiating spermatogonia in mice (A1-4, Intermediate, B); four in monkeys (B1-4) and one in humans (B).

### Consensus SPG6

In SPG6, all species turn off DMRT1 while maintaining DMRTB1 expression (not annotated in macaque), which is a specific signature of mouse Type B spermatogonia^12,30^. In addition, SPG6, all species begin to express many markers previously identified in mouse preleptotene spermatocytes including Tex101, Ly6k and many established meiotic genes such as Sycp2/3, Syce1/2,Meiob, Prdm9, Spata22, and DNAJB11. Therefore, SPG6 marks a transition from type B spermatogonia to early meiotic preleptotene cells in all three species. However, we identify many new genes that are either expressed in all three species (i.e. FMR1NB, ZCWPW1, DPEP3, IQBP1, CALR), in primates only (BEND2, ZRANB2, PAGE4, ZNFX1, HLTF, ZBED5) or singularly expressed in mouse (Swt1, Tgfbr1, Rhox13, PhTF1, Fmr1), macaque (ZNF850, MAGEB17, ZMYM1, TCEA1) and human (PRDM7, full VCX family, ERBB3, ERVK13-1, SSX3, TRIML2). While functional significance for many of these species-specific genes in meiosis remain unknown, these genes may provide new insights into the evolution of meiotic entry program (see **Table S3**A for comprehensive list of markers that are shared or species unique genes).

Nomenclature that has evolved to describe spermatogenic lineage development in rodents is different than monkeys or humans, which makes it difficult to translate the implications of data produced in one species to the other species. Therefore, in an effort to unify our transcriptionally distinct populations with classical morphometric descriptors, we updated a model comparing spermatogenic lineage development across species **Figure S2D**^9^. Briefly, **SPG1** and **SPG2** represent two distinct populations of undifferentiated spermatogonia which do not strictly segregate with classic descriptions of A_dark_ and A_pale_ spermatogonia but are comparable to the mouse undifferentiated spermatogonia populations (A_s_-A_paired_-A_Aligned_) (see below). **SPG3** contains cells that are transitioning from undifferentiated (A_al_ mouse; A_d/p_ monkey and human) to differentiated (A1 mouse; B1 macaque; B human). **SPG4** and **SPG5** are transcriptionally distinct sub-populations of transit-amplifying differentiating spermatogonia (A2-4, Intermediate mouse, and Type B; B2-B4 monkey and B human). **SPG6** cells have upregulated the meiotic program and represent Type B to preleptotene spermatocytes in all three species.

Altogether, our analysis identifies six molecularly defined SPG states which are comparable across species, but not based on a few preselected marker genes. The equivalency of the SPG states across species is confirmed by the average cell-cell Jaccard distances between clusters (**Figure 2F**). On a global scale, cluster-cluster similarities between species are weaker in earlier SPG states when compared to SPG5-6 across species, suggesting that the start states and molecular pathways for early spermatogonia differentiation are variable across species, but these cells ultimately reach a more conserved molecular identity, or experience a convergence in differentiation program prior to meiosis. Our analysis also reveals both similarities and differences between species in marker gene identity and timing of expression (summarized **Figure 2G**). While previous studies assumed that well known markers are transferable across species, we now provide a foundational set of universal and species-specific markers that can be used to isolate and analyze equivalent cellular states in human, monkey and mouse.

### Molecular states SPG1 and SPG2 do not correlate with A_dark_ and A_pale_ histological states in human

Historically, the undifferentiated spermatogonia in humans and macaques has been described by intensity of nuclear staining with PAS hematoxylin as A_dark_ and A_pale_ spermatogonia^5,31,32^. Based on in vivo mitotic labeling with^3^H-thymidine, Clermont proposed that A_dark_ were quiescent (reserve) stem cells and A_pale_ are active stem cells that maintain steady state spermatogenesis^7^. To determine whether SPG1 corresponds to either histological state, we combined PAS and hematoxylin staining with immunohistochemistry of human testis cross sections for SPG1 markers PIWIL4, MORC1 and TCF3. PIWIL4 and MORC1 are enriched in SPG1, while TCF3 is more equally distribute between SPG1 and SPG2 (**Figure 2E-F**). All three markers were localized to the basement membrane of human seminiferous tubules and were expressed by both A_dark_ and A_pale_ spermatogonia (**Figure 3E-H**). However, TCF3, which is expressed by cells in both SPG1 and SPG2, was skewed more toward A_pale_ spermatogonia (**Figure 3G**). Thus, SPG1 contains both A_dark_ and A_pale_ spermatogonia, indicating that these descriptions of nuclear morphology do not necessarily segregate transcriptionally. However, there may be a subpopulation of TCF3+ A_pale_ with a distinct transcriptional program (SPG2).

### Human spermatogonial stem cell activity is enriched in, but not exclusive to SPG1

While it is tempting to predict stem cell potential for a population based on developmental ordering of single cell data or expression of markers, the only way to truly test spermatogonial stem cell (SSC) potential is by transplanting cells into autologous/homologous recipient testis to reconstitute spermatogenesis^33^. Autologous/homologous transplantation in humans for experimentation is not possible. For humans, the gold standard is xenotransplantation into the testes of busulfan-treated immunodeficient mice (NCr nu/nu; Taconic)^34,35^. Although hSSCs do not complete spermatogenesis in this context, they do migrate and colonize the basement membrane of the seminiferous tubule, where they proliferate and produce characteristic chains of spermatogonia that survive for months (**Figure 3I**).

In order to functionally characterize the colonization potential of the SPG1 population, we enriched for these cells by FACS using the SPG1 cell surface marker TSPAN33 (**Figure 2E, Figure 3J**) that was also described in the human undifferentiated spermatogonia clusters of two previous studies^22,23^. Unsorted, TSPAN33-negative, and TSPAN33-positive cells were xenotransplanted into recipient nude mice (**Figure 3I-J**). Analysis of recipient testes two months after transplantation revealed that stem cell concentration was significantly enriched in the TSPAN33 fraction (46 ± 15 colonies/10^5^ cells transplanted) compared with the Unsorted (10.7 ± 3 colonies/10^5^ cells) and TSPAN33-negative fraction (3.2 ± 0.9 colonies/10^5^ cells) (**Figure 3K**). This confirms that the TSPAN33+ population (merged SPG1) is enriched for transplantable human spermatogonia, but it does not capture the entire population. In fact, when adjusting for the total number of cells in each fraction, it becomes clear that almost half of colonizing activity resides outside of the TSPAN33+ fraction (Figure 3L). Thus, transplantable human spermatogonia are transcriptionally heterogeneous, at least with respect to TSPAN33 expression. Future studies using combinatorial staining, sorting, and transplantation experiments will be needed to determine the phenotype(s) of TSPAN33-negative human SSCs. Molecular heterogeneity in the pool of undifferentiated stem/progenitor spermatogonia may allow spermatogonia to respond to continuously changing niche signals and maintain tissue homeostasis, as suggested by Guo and colleagues^22^.

### Generation of a universal pseudo-timescale for mammalian spermatogenesis

The time required to complete a full spermatogenic cycle is variable across species (~35d in mouse^36,37^ vs. ~42d in macaque^38–40^ vs. ~74d in human^37,41^). Histological analysis of pulse labelling and cell ablation have been used to determine the length of major spermatogenic processes, but to what extent germ cell trajectories and molecular states align across species is poorly understood. To address this gap, we used our scRNA-seq data to reveal heterochrony in the spermatogenesis process and generate a universal spermatogenesis pseudotime comprised of aligned cellular states across species. which will allow us for the first time to define conserved and diverged cellular and molecular processes.

To align the three species, we collapsed the spermatogonia populations to a single group, and ordered the remainder of non-SPG germ cells (N=18537 mouse, N=12016 macaque, N=4293 human) along a linear trajectory using 2,000 highly expressed and highly variable genes selected independently for each species. We then defined distinct pseudotime-points by using *Monocle2*, which divided cells into 200 ordered “bins”, each corresponding to a biological state. To compare across pseudotimes between species, we calculated a rank correlation of each of the 200 states between each species pair, using only a shared subset of highly variable 1:1:1 orthologs detected in all three species (highly variable genes/species = 2000, Shared N=247 genes) (**Figure 4A**). Therefore, if the differentiation process started and ended in equivalent states between two species and advanced at the same pace through intervening states, a perfect diagonal line would be observed. However, heterochrony occurs between species as deviations from the diagonal are observed (**Figure 4A**), suggesting that the biological steps defined separately for each species - using the current standard method - proceed at different rates, even when the comparison is based on the same set of 247 genes. We subsequently confirmed that this pseudotime-warping pattern is not driven by the use of 247 common genes in the initial cell assignment, because when we reassign cells to an alternative set of 200 bins, defined for each species separately by using their own highly variable 2000 genes (**see Methods**), there is little difference in the between-species alignment patterns (**Figure S4A**).

**Figure 4.**
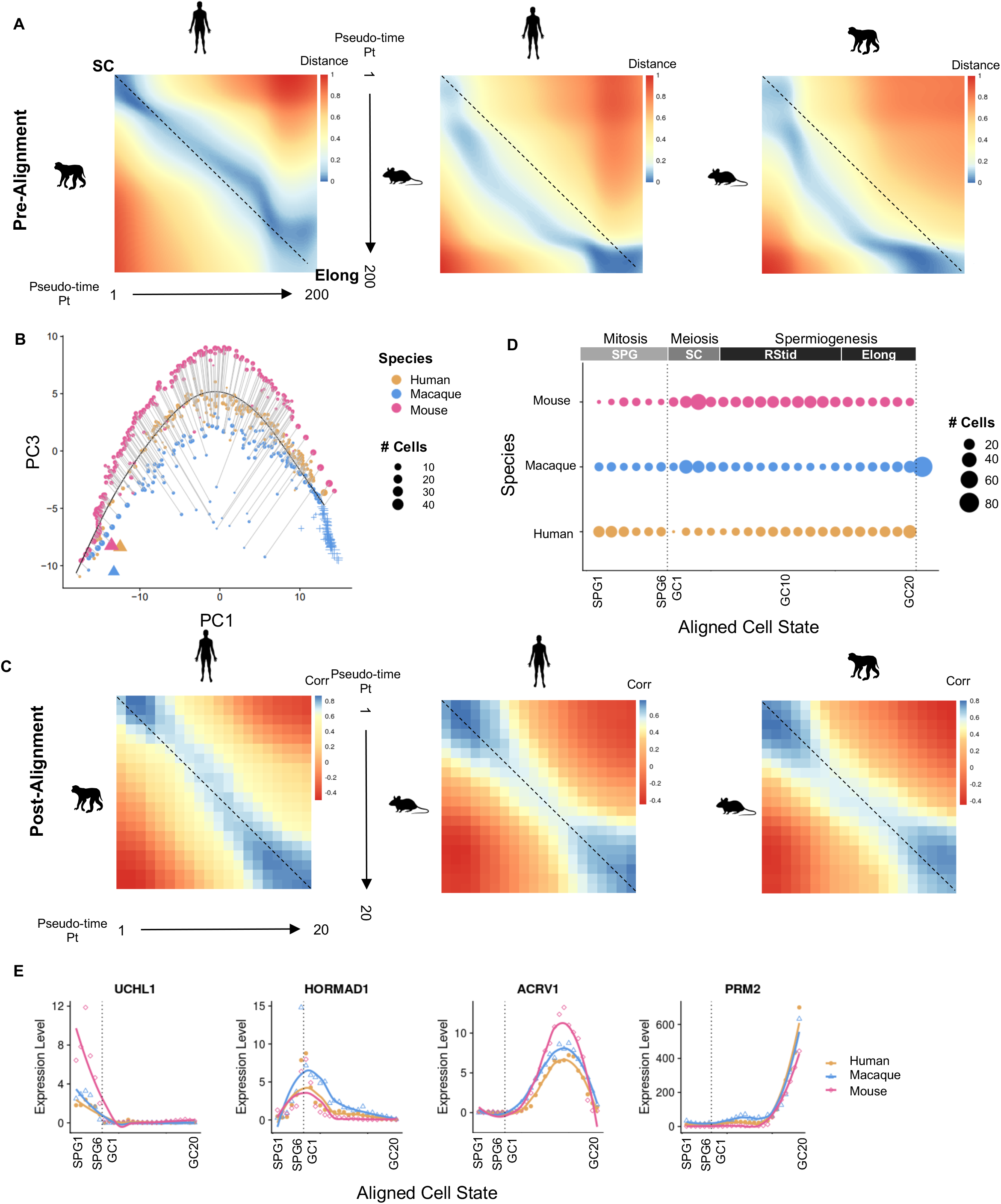

Although the Human and macaque pseudotimes were largely aligned, the final ~1/3 of the macaque trajectory maps to the final ~1/5 of the human trajectory, suggesting that most mature or terminal state (elongating spermatids) in human corresponded biologically to a wider span of the mature states in macaque (**Figure 4A, left panel**). Furthermore, we found that the last ~10% of molecular states in macaque appear to extend further beyond the final states of either human or mouse, suggesting that macaque spermatids progress molecularly further than humans or mouse (**Figure 4B**). Targeted analysis of these overhanging states in macaque revealed no unique markers (data not shown), but rather progressive loss of pre-existing transcripts regardless of UMI depth **(Figure S4B-C**). This observation is consistent with the previously reported RNA degradation in human sperm and mouse sperm^42–44^. Whether these differences are reflected in the transcript content of mature sperm or paternal contribution of transcripts to the zygote remains to be determined.

As expected, more distantly related species showed greater mis-alignments of their pseudotime (**Figure 4A**). For example, human-mouse and macaque-mouse comparisons showed heterochrony at both the start and the end of the trajectories (**Figure 4A, middle, right panels**). To directly compare equivalent molecular states across human, macaque, and mouse we developed a common pseudotime shared by the three species by fitting the three individual species pseudotime-series to a single principle curve in PC1-PC3 space (**see Methods**) (**Figure 4B**). Cells were assigned to the nearest point on this consensus trajectory, now divided to 20 bins. The final states of the macaque trajectory that did not have an equivalent state in human or mouse (**Figure 4B**), were added as a single group at the end of the aligned 20-bin trajectory, representing the 21st bin in macaque (**Figure 4D**). As expected, the 20 centroids, obtained for each of the three-species using the universal coordinates now showed a straight diagonal in the 20-by-20 distance matrix (**Figure 4C**), confirming their synchrony after warping to the universal time.

With analogous states properly matched, we compared cell occupancy within the 20 (mouse and human) or 21 (macaque) post-mitotic germ cell differentiation states, along with the six spermatogonia states defined above for each species (**Figure 4D**). Based on the temporal profiles of several conserved marker genes (**Figure 4E**), we mapped the 26 (27 in macaque) molecular states in each species to major spermatogenesis processes (**Figure 4D**): mitosis (6 spermatogonia bins as defined in the SPG section above), meiosis (~bins 7-11), and spermiogenesis (~bins 12-20 round spermatids and ~21-26 elongating spermatids). Macaque has an extra bin (the 27th) accounting for its unique terminal state in elongating spermatids. The number of cells per bin can reflect the rate of traversing developmental states and/or the balance of cell division and death. For example, humans and macaques, unlike mice, have a larger pool of undifferentiated spermatogonia (**bins 1 and 2; Figure 4C**) that undergo a limited number of transit amplifying divisions, whereas, mice have a much smaller population of undifferentiated spermatogonia that undergo a series of mitotic amplification divisions (**bins 1 and 2; Figure 4C**) (see Spermatogonia section). The cell numbers in bins 5 and 6 (SPG populations) in macaque and human were maintained relatively constant, but a reduction was observed in mouse, consistent with a theoretical 50% reduction in spermatogonia estimated to occur *in vivo*^3,45^. Furthermore, in bins 7-10, which represent the transition to meiosis and meiotic divisions, the mouse and macaque states expectedly display an expansion of cells, whereas, humans appeared to initially face a significant bottle-neck at the transition to meiosis. However, only a portion of this reduction can be accounted for by previous histological estimations of degeneration rates in humans^46^, suggesting that human germ cells may uniquely proceed more quickly through initial stages of meiosis. Finally, we find a greater number of cells towards the end of the trajectory in primates, likely reflecting molecular differences in spermatid differentiation, as there are no cell divisions occurring post-meiotically. Thus, our cross-species alignement of spermatogenesis supports previous histological estimations but also identifies divergence in the amount of molecular states allocated to certain processes, suggesting unique regulation.

In summary, our dataset establishes the first molecular coordinate system for the temporal progression of male germ cell differentiation for three mammalian species and provides markers for individual steps for each species (**Table S4**) which will be a valuable resource for the community for future experiments.

### Species-specific changes in gene expression across the germ cell developmental trajectory

To identify modules of genes with distinct phases of activation in each species, we employed k-means clustering of 2000 highly variable 1-1-1 orthologs expressed across aligned pseudo-time points independently in mouse, human, and macaque (**Figure 5A**). This unbiased gene clustering analysis revealed successive waves of gene expression corresponding to major biological processes of mitosis (K=1), meiosis (K2-3), and spermiogenesis (K4-6) confirmed by gene ontology (**Table S5**). To determine if genes were activated at the same or different phases across species, we used the union of highly variable 1-1-1 orthologs in each of the species pair, ordered them by the six K-means gene group designations, and linked the orthologous genes by lines (**Figure 5A**). Genes expressed “in phase” (I.e. in K1 of both human and macaque) in two species are linked by horizontal lines and are considered to have conserved temporal expression patterns, whereas, genes that have “shifted” expression patterns are not temporally conserved and are linked by diagonal lines. Finally, genes without a connecting line are those that have “unique” expression patterns, such that their ortholog is not detectable or not dynamically expressed in the other species, and thus cannot be assigned to a K-means group.

**Figure 5.**
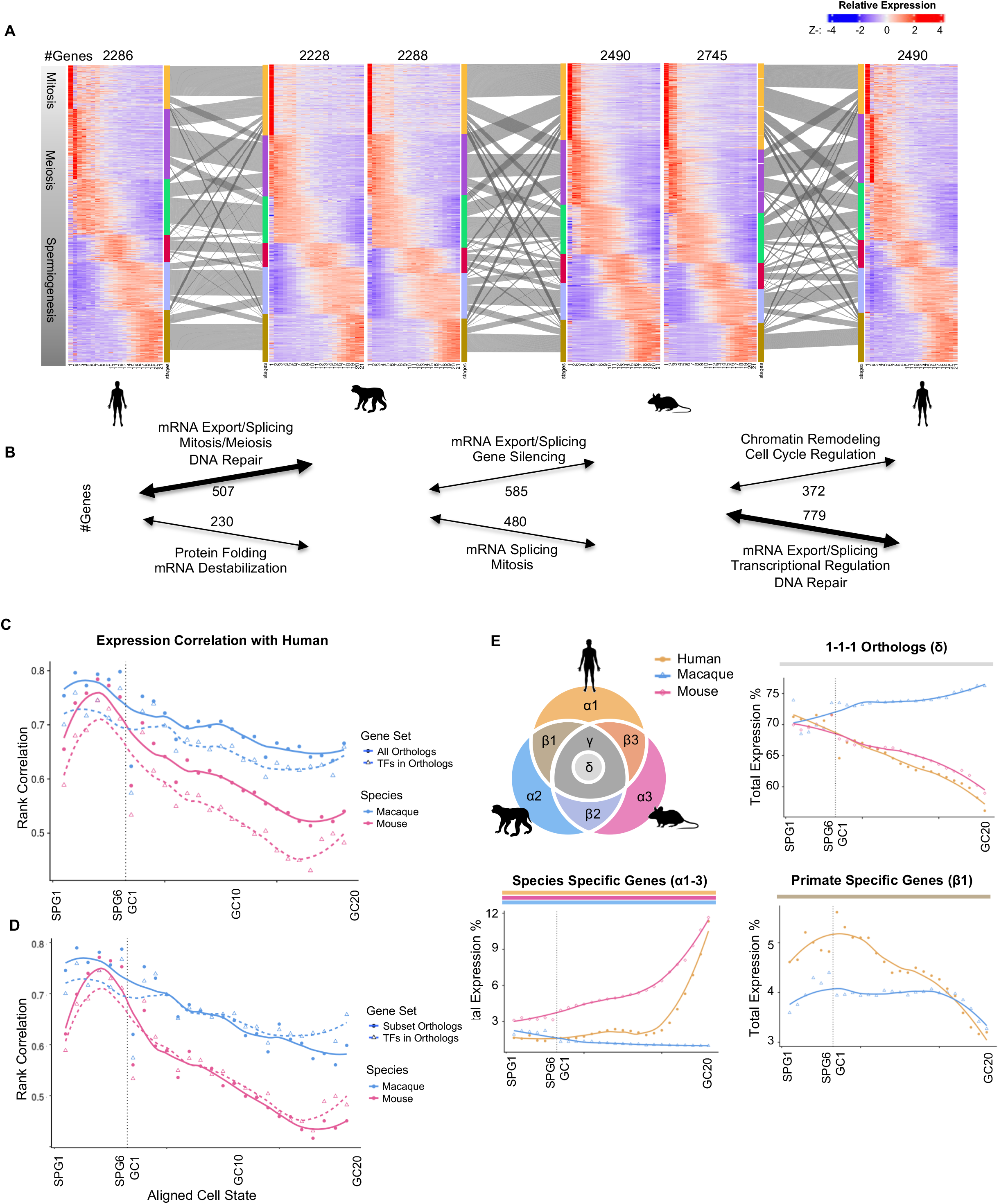

As expected, more closely related species displayed higher phase conservation (**Figure 5A, S5A**). For instance, a comparison of human-macaque showed more in-phase genes (40-56%) than phase-shifted (25-38%), whereas human-mouse showed the opposite trend (20-38% in phase vs. 31-48% shifted), as did macaque-mouse (26-45% vs. 36-49%) (**Figure S5A**). The shifted genes may reflect changes in the functionality or regulation of genes over the course of evolution. The shifted genes were found to be involved in diverse biological processes such as DNA repair, RNA splicing/processing/binding, and transcription regulation. The direction of shift, revealed some separation of functionality (**Figure 5B**). For example, human genes that lagged in expression compared to either macaque or mouse, included numerous genes involved in mRNA export and splicing including genes such as RNPC3, NONO, PRPF8, and SNRPB. Broadly, many RNA-binding proteins exhibit shifts in expression across species, including deadbox genes, an evolutionarily conserved family of RNA-helicases. DDX4 (Vasa) is specifically expressed in germ cells from fruit flies to mammals^47^, whereas many additional members of this family are also expressed during spermatogenesis with greater variation among species. For example, while DDX18 and DDX46 were expressed early in K1 in humans, these genes were not expressed until K2 and 3, respectively in mouse. Reciprocally, DDX23 is expressed early (K1) in mouse, but later in human and macaque (K2), while yet another member, DDX52 is expressed meiotically in (K2) in mouse, but significantly later in human spermiogenesis in K5. The precise functions of these DDX family members have not been determined, but their expression patterns illustrate potential diversification of functions in spermatogenesis. Future analyses are needed to determine whether the increase in out-of-phase genes reflects overall reduced conservation of gene expression during evolution, or programmatic differences in the regulation of spermatogenesis.

Additionally, we find that the number of genes with unique species-specific expression increased progressively across the different k-means groups across all pairwise comparisons. For example, the human vs. macaque, K=1 gene group has 10% unique genes whereas K=6 has ~25% unique (**Figure S5A**). Therefore, this analysis identifies numerous species-specific transcripts (unique or shifted transcripts) that may be important for the germ cell developmental program.

### Overall divergence in expression increases across spermatogenesis in mammals

While it is known that testis is one of the most rapidly evolving tissues on both a sequence level and transcriptome level^48^, our study offers a new opportunity to compare the degree of gene expression conservation across different stages of spermatogenesis. Using 1-1-1orthologs we calculated the rank correlation between gene expression centroids for pairs of species, for each of the 26 ordered stages: 6 for SPG and 20 for the germ cells. The between-species correlation is highest in spermatogonia and decreases progressively through differentiation to spermatids (**Figure 5C, solid lines**). The correlation values for 835 transcription factors (TF) showed similar decreasing patterns, albeit with lower levels than for all orthologs (**Figure 5C, dotted lines**). To assess if the lower correlations for TFs than for all orthologs are partly due to the lower expression levels of TFs, we re-calculated the correlations for randomly subsampled orthologs of the same number (835) and matched gene expression distribution as the TFs (**Figure 5D**). After this resampling-based correction, the TF correlations are no longer lower for meiotic and post-meiotic germ cell states but remained lower than the matched orthologs in SPG states. This result suggests a greater between-species programmatic divergence for transcription factors than for non-TF genes, and this effect is mainly observed in SPG stages.

Many TFs may be expressed at low levels in scRNA-seq data, yet their functional effects can be estimated by the concerted changes of their downstream target genes, i.e., each TF’s “regulon”. By using *SCENIC*^49^ we obtained the activity scores for 82 and 88 regulons with 10+ target genes for human and mouse germ cell centroids, respectively (Table X). We ranked their degree of conservation by the human-mouse correlation of regulon scores across the 26 stages, and summarized the top 17 most conserved and top 5 most diverged between human and mouse (**Fig S5C**). (This analysis was not performed in macaque since a corresponding TF database is not available). Importantly, many of the top TF regulons (e.g. EZH2, E2F1, YY1 and Sox6) we identify have been previously implicated in spermatogenesis^50–54^. Second, the phasic pattern we report here is consistent with previously described functional dynamics in mice^50–54^. However, our results also include several novel regulons that have not been previously explored in the context of spermatogenesis (e.g., UBTF, GABPA, BRG1, GABPB1, BACH1) and may play a role in germ cell epigenetic memory or transcriptional regulation.

While this comparison takes into account only orthologous genes, it is possible that other species-specific genes may have temporal bias in expression and thus contribute to observed divergence in the spermatogenesis program. Next we examined the bulk gene expression levels for genes of various ortholog classes (**Fig 5E, S5B**). Macaque appeared largely unchanged in the amount of 1-1-1 ortholog expression, and macaque-specific genes contributed little to total expression, likely a reflection of low annotation quality of the macaque genome where the vast majority of genes are orthologs (**Figure 5E, S5B**). This is further underscored by the high number of orthologs overlapping in human and mouse only (**Fig S5B**), many of which likely have currently unannotated genes in macaque. Focusing instead on categories with greater annotation confidence, several interesting patterns are observed. Human and mouse show a decrease in the degree of ortholog expression across spermatogenesis, which is complemented by an increase in species specific (lacking an ortholog in mouse or macaque) gene expression (**Fig 5E**). The amount of primate specific gene expression (present exclusively in both human and macaque, regardless of copy number) peaks at the entry into meiosis. Interestingly, these primate specific genes contain a large number of C2H2 zinc-finger containing proteins (227/875 human orthologs), about half of which have been characterized as having testis-enriched expression relative to all other tissues in the Human Protein Atlas. Most of these genes (198) are KRAB domain-containing, the largest subgroup of C2H2 zinc-finger proteins which have been shown to bind endogenous retro elements (ERE)^55^. Indeed many of these genes are capable of binding at least one ERE in vitro, with most binding ERVs (**Figure S5D,E**).These genes may then aid in identifying novel primate-specific regulators and mechanisms of entry into meiosis.

### Species specific transcriptomic differences in interstitial somatic cells suggest functional divergence between rodents and primates

Germ cell maturation requires intricate communication between the germline and soma. The importance of germline-soma compatibility has been elegantly underscored *in vivo* by cross-species transplantation experiments, where rat SSCs can complete spermatogenesis in the mouse testis^56^, whereas macaque or human SSC colonize the mouse testis but fail to initiate meiosis^57^. The incompatibility between SSCs and somatic compartments of distant organisms may reflect changes in signaling pathways or cell types in the microenvironment that alter regulatory programs within SSCs.

To create a new atlas of the somatic compartment we performed focused re-clustering of 3,722 human and 2,098 macaque somatic cells, yielding 7 clusters for each (**Figure S6A-B**). Using well-established markers, we identified myoid cells, endothelial cells, Leydig cell precursors, and macrophages, and several rarer cell types not extensively studied in testis, such as T cells and two types of pericytes (**Figure 6A-B, S6C**). However, unlike mouse our human and macaque datasets failed to capture Sertoli cells, Innate Lymphoid cells, or mature Leydig cells^12^, likely due to the mechanical or physiological stress of the cryopreservation and dissociation methods.

**Figure 6.**
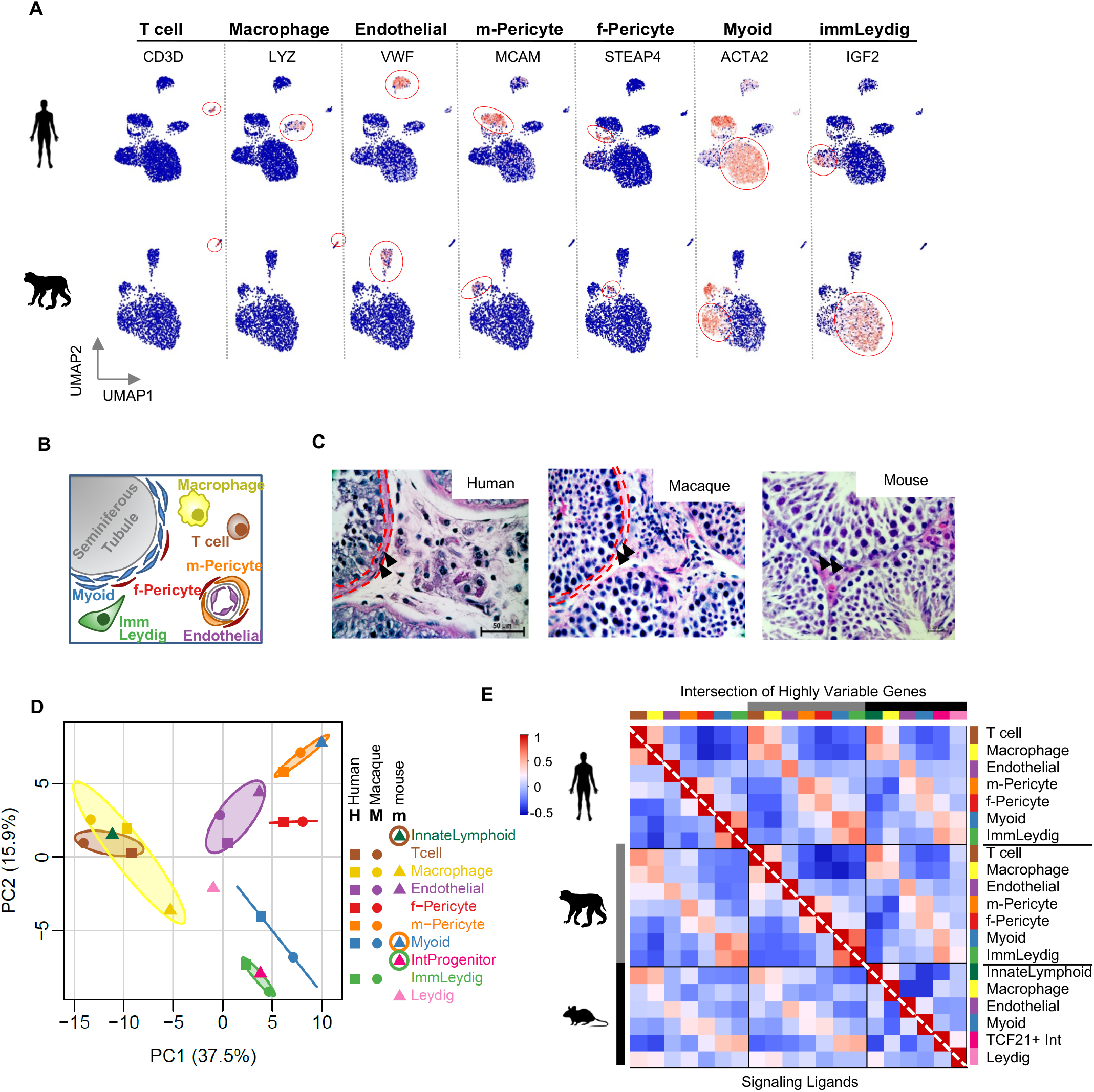

One of the most abundant populations captured in primates was peritubular myoid cells, defined by the expression of MYH11, ACTA2, collagens, laminins, versican (VCAN), and fibronectin (**Figure 1C, 6A, S6C, Table S6**). This overrepresentation of myoid cells in primates, as compared to mouse^12^, reflects structural difference in seminiferous tubule basement membrane, which consists of multiple layers of myoid cells in human/macaque, but a single layer in mouse (reviewed in^58^) (**Fig 6C**). Myoid cells are contractile and are involved in the transport of spermatozoa and testicular fluid in the tubule^59^. Recent studies in mice have suggested that myoid cells can also secrete a number of substances, including extracellular matrix components (*in vitro*) or growth factors that may modulate the function of neighboring somatic cells (i.e. TGFB or Activin A) or spermatogonial cell proliferation (i.e. GDNF, *in vivo*). Consistent with these findings, human and macaque myoid cells expressed many genes involved in muscle contraction, as well as, some conserved (fibronectin and collagens) or unique extracellular matrix (ECM) proteins such as decorin and biglycan (**Table S6**), which are believed to be absent from the mouse testis under normal cellular homeostasis^60^. Unlike mouse myoid cells, the human and macaque myoid cells don’t produce the spermatogonial stem cell proliferative factor GDNF, but expressed components of other signaling pathways like PDGFRA, EGFR, PTCH1, and PPARA which are not detected in the analogous mouse population (**Supplemental Table 6**). Therefore, although both mouse and primate myoid cells have contractile properties, they have acquired species-specific signaling and molecular functions.

A second major population identified in the human and macaque is immature Leydig cells, which is observed in neonatal rodent testis, but thought to be absent in adult mouse testis (reviewed in^61,62^) and was lacking in our adult mouse testis data^12^. This human and macaque population expressed multiple markers previously shown to be restricted to spindle-shaped putative progenitor Leydig cells^63^ and are critical for Leydig cell differentiation^64,65^ (e.g. DLK1 (unannotated in macaque), IGF1/2, CFD (Adipsin), SFRP, PTCH2, and IGFB3 (**Figure 6A, Figure S6C, Table S6**)), whereas their expression of steroidogenic enzymes (STAR and Cytochrome P450 genes) were very low and limited to a small subset of cells (data not shown). These cells also expressed ECM genes such as decorin and collagens. Additionally, we found that the immature Leydig cell transcriptome is highly similar to that of myoid cells (**Figure S6B**), suggesting that myoid and Leydig cell lineages may share a common progenitor in human and macaque, as seen in mice^66,67^.

The somatic compartments also contained an endothelial cell (VWF, NOSTRIN, AQP1, CD34 (human)) population and a heterogeneous population of cells known as pericytes, which together interact to form the mural wall of small blood vessels throughout tissues^68,69^. Pericytes express MCAM, RGS5, PDGFRB, NG2, CCL2, ABCC9, and NOTCH3, consistent with recently defined markers in brain and lung datasets^70,71^ (**Fig 6A, S6C, Supplemental Table 6**). Furthermore, the testis pericyte population splits into two distinct sub-clusters of pericytes that can be uniquely separated into a muscular (m-pericytes) or fibroblastic (f-pericytes) types based on their expression profiles (m-pericytes: MCAM (higher expression), CRIP1/2, RERGL, ADIRF (unannotated in macaque), and several myosins MYL9, PTM1/2) (f-pericytes: STEAP4, GUCY1A1/2, ITGA1, CD36 (human), CD44(human) and several collagens and laminins) (**Figure 6B, S6C, Table S6**). Using immunohistochemistry data available from the Human Protein Atlas, we find that several markers of m-pericytes (MCAM, CRIP1, ADIRF) are more restricted to cells surrounding blood vessels; whereas f-pericyte markers (CD36, CD44, ITGA1) are present both around blood vessels and in the interstitial space (**Figure S6D**). Despite the high correlation between the two pericyte populations (**Figure S6B**), the individual species cluster-cluster centroid correlations suggested that the transcriptome of m-pericytes was more highly correlated to myoid and immature Leydig cells, whereas f-pericytes were more distinct (**Figure S6B**). These findings suggest that f- and m-pericytes might arise from different cells of origin in the testis.

Finally, we identified several types of immune cells. The majority were macrophages (CD163, HLA-DRA/MAMU-DRA, LYZ, TYROBP) (**Figure 6B, S6C, Table S6**). Although rare, lymphocytes were also present in the testis. In rodents, CD8+ (cytotoxic) T cells are the predominant lymphocyte population^72^, but in humans, the frequency and location of lymphocytes is poorly understood^73–75^. Here we identified T cells (CD3D, TRAC, TRBC2, CD69) (**Figure 6B, S6C, Table S6**) with some CD8 expression (23% of T cells in human, 10% in macaque), indicating that CD8+lymphocytes are indeed present in adult human and macaque testis.

Taken together, we identified many of the expected major or rarer somatic cell populations of the testis that express shared or species-specific factors. Given the several noted differences in the marker genes of primate vs. mouse somatic cells, we sought to compare somatic cells more globally to uncover how these somatic populations have changed over the course of mammalian evolution. Briefly, we merged the somatic cells from all three species and performed principal component analysis (**Figure 6D**) and rank correlation of the cell type cluster centroids (**Figure 6E**) using 1-1-1 orthologs (**see Methods**). In the merged PCA plot, PC1 and PC2 segregated the immune cell populations (macrophages and Tcell/ILCII) from the remaining somatic cells, confirming that the widest distinction is driven by the rapid evolution of cell type differences, rather than species differences (**Figure 6D**). Although, we did not previously identify T cells in mice, mouse innate lymphoid cells had the highest correlation with primate T cells, as expected based on their similar origin from a common lymphoid progenitor and shared functions^76^. Endothelial and Macrophage cell transcriptional programs appeared to be largely conserved as they have higher correlation values across species (r=0.6-0.8; **Figure 6E, upper half of matrix**).

However, there are notable between-species differences in the remaining somatic cells. For example, mouse myoid cells appeared to be more transcriptionally similar to primate m-pericytes, than primate myoid cells. Similarly, although Leydig precursor cells in primates somewhat resembled adult mouse Leydig cells, they were more similar to our previously unexpected mesenchymal cell population in the mouse testis^12^. Thus, many of the somatic cells within the testis have morphed transcriptionally across species. To confirm that the unexpected cellular associations are not driven by orthologous gene selection, we performed clustering using the union of marker (**Figure S6E**; **lower half of matrix**) or the highly variable genes from each of the three species (**Figure S6E**; **upper half of matrix**), and show similar cellular association and correlation patterns. Using ~300 genes encoding known ligands, we found a nearly identical pattern of correlation among the cell types and across species (**Figure 6E, lower half of matrix**), indicating that the observed shifts in cell identity relationships is likely due to genuine global programmatic changes in the transcriptome, signaling pathways and possibly function.

Finally, as expected, the strength of the cell type correlations across species decreases with increasing evolutionary distance. How these changes impact spermatogenesis are unclear, but they may contribute to the incompatibility observed between germ cells and somatic cells of distant species.

### Evolutionary differences in the germ cell - soma cell communication

Our scRNA-seq data also provides an unprecedented opportunity to begin exploring the poorly understood cell-to-cell communication between the soma and the germline in the testis. To comprehensively catalogue the cell-to-cell communication we filtered known ligand–receptor interaction pairs in public interaction databases^77^ by focusing on those with 1) ligands detected in somatic cell clusters and receptors detected in germ cell clusters, and 2) either the ligand and receptor were highly variable in at least one species. This led to 1059, 529, and 785 L-R pairs in human, monkey, and mouse, respectively (**Table S7**). By calculating the Interaction Scores for each L-R pair (**Methods**) across all 182 cell type pairs (7 somatic and 26 germ cell types), we created a somatic-to-germ cell interaction map that depicts the number of putative strong interactions (defined in Methods) for each cell pairs (**Figure 7A**). This global analysis reveals that many somatic cells do have the potential (i.e., the expression of ligands) to communicate with germ cells, supporting an open niche model for the mammalian testis. In humans and monkeys, the immature Leydig cells, Myoid and f-pericytes have the most extensive putative interactions than other somatic cell types detected in our dataset. Meanwhile, Sertoli and Leydig cells – two central niche cell populations - have the least number of outgoing signaling interactions in mouse. Instead, the Tcf21+ interstitial population, myoid, macrophage, and endothelial cells exhibit the greatest number of interactions (**Figure 7A**). Among the germ cells, the spermatogonia populations (SPG1-6) tend to have more putative L-R interactions than the germ cells, consistent with the role of the blood-testis barrier (BTB) that prevents meiotic and post meiotic germ cells from directly communicating with neighboring somatic cells. A possible exception may be when the BTB breaks (stages VIII-IX), giving differentiating germ cell populations the opportunity to communicate with surrounding somatic cells and possibly explaining the weak and few interactions detected between somatic cells and later germ cell stages.

**Figure 7.**
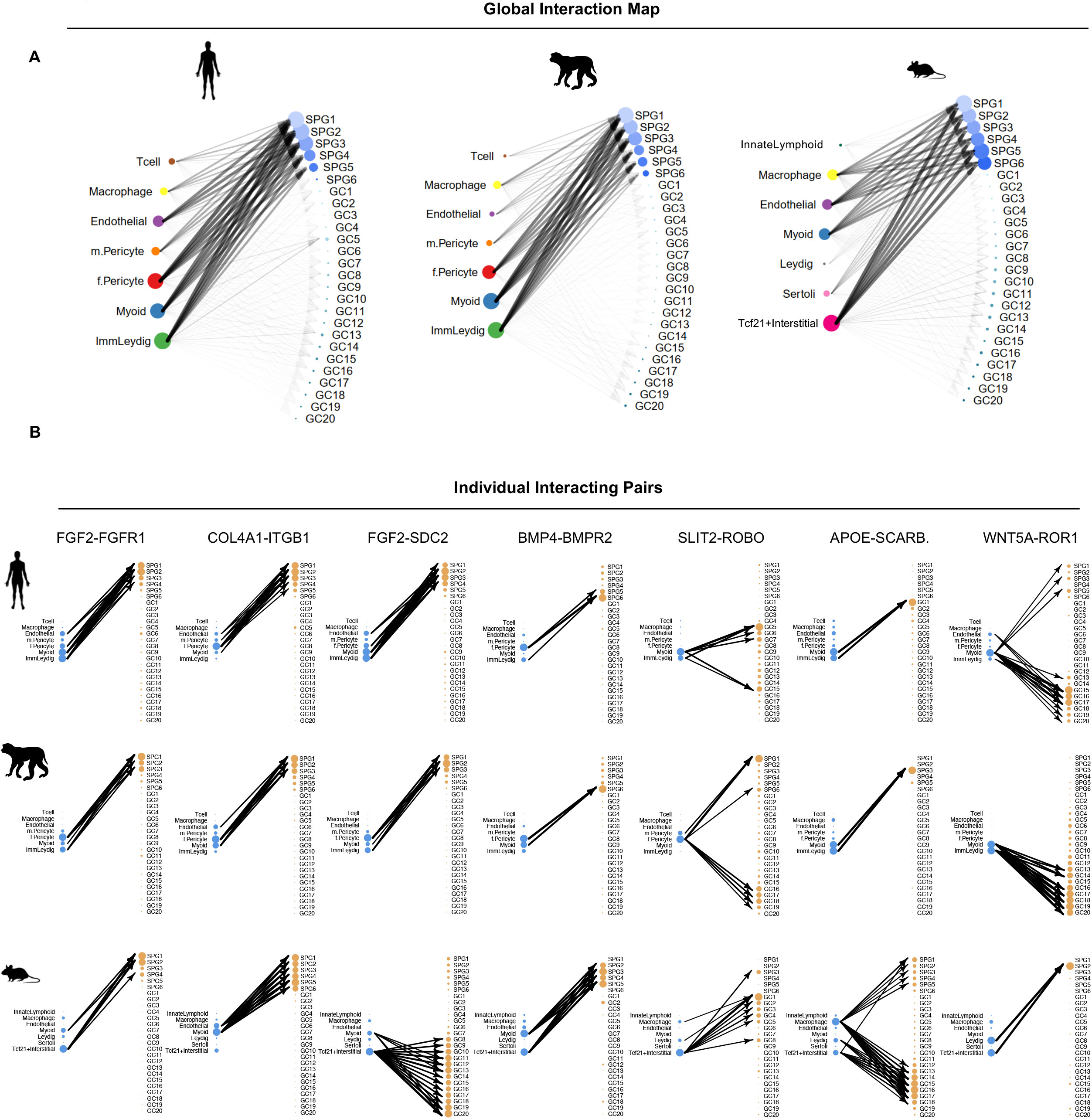

Given the significant signaling differences observed across cell types, we explored the conservation of individual signaling pairs. For each L-R pair we calculated its between-species correlations across the 182 cell type pairs and used such a measure to rank the degree of conservation of signaling pathways (see **Table S7** for the full list). Globally, **Fig S7A-B** shows the 30 most conserved and 30 least conserved L-R pairs, from which we highlight examples for several classes of conservation patterns (**Fig 7B, S7C**). First, the target cell is conserved but the source cell is not (FGF2-FGFR1, COL4A1-ITGB1, WNT5A-RYK; CXCL12-ITGB1), while for some pairs there is better conservation within primates specifically (FGF2-SDC2; BMP4-BMPR2). Second, the source cell of the ligand is conserved but the target cell is not (FGF2-SDC2). Third, both the source and target cell type vary across species (SLIT2-ROBO; APOE-SCARB1; WNT5A-ROR1; BMP4-BMPR1B; FGF2-SDC4; JAG1-NOTCH2; FGF7-FGFR3). These patterns suggest that while some pathways may be employed in multiple mammalian species to regulate similar stages of spermatogenesis, the timing of activation and the origin or target of the signal may have diverged. In sum, we leveraged ligand and receptor expression levels in specific cell types to generate testable hypotheses regarding intercellular signaling patterns and potential evolutionary shifts. Future perturbation and spatial transcriptomics and proteomics experiments will be needed to validate them.

## Discussion

Germ cell differentiation is a highly regulated process reliant on both intrinsic and extrinsic signals. Although the basic cellular events - mitosis, meiosis, spermiogenesis- are largely conserved across species, there is notable divergence in the cellular dynamics, structural organization, hierarchical patterns, and output of this differentiation process^9,11^. Molecular and genetic analyses of mouse testes have provided substantial insights into the process of spermatogenesis and are often used to guide parallel studies in primate testes. However, most comparative studies to date are reliant on a handful of markers developed in histological analyses, which are used to define presumably equivalent cellular states across species. These approaches may overlook important functional divergence between species, and may yield an incomplete, or even incorrect, picture of primate spermatogenesis. Therefore, an unbiased comparative analysis of germ cell development and somatic microenvironment is needed to elucidate similarities and differences across mammalian spermatogenesis. We addressed this challenge by employing single-cell RNA-seq analysis of normal adult human and macaque testes, and comparing with our previous mouse dataset to provide a new generation of cellular atlases within each species and across species

The first contribution of this study is the complete differentiation trajectory of the spermatogenic process in human and macaque, along with molecular markers for major germ cell and somatic cell types, which we provide separately for each species (**Table S1**). On a global scale, we observe instances of inter-species difference in transcription regulation, suggesting evolutionary changes of some aspects of spermatogenesis program. For example, we find that meiotic sex chromosome inactivation (MSCI) is incomplete in primates; we estimate at least 16 genes in human and 14 genes in macaque escape sex chromosome repression in spermatocytes. Several of the identified escapees appear to be critical for normal germ cell development, e.g. DYNLT3 is involved in meiotic chromosome segregation^78,79^, while AKAP14, AKAP4, CYLC1, and PIH1D3 are important for sperm structure and motility^80–85^. Several escapees (SPANXN3, SPANXN5, SPANXD) are members of the primate ampliconic SPANX gene family^86^. Although the function of these genes is unknown, copy number variants involving SPANXN5 have been linked to human infertility^87–89^, indicating this family may have critical roles in spermatogenesis.

To directly compare germ cell states, we combined and re-clustered spermatogonia from all three species, allowing us to identify six consensus states. While different vocabulary has been developed to describe spermatogonial development in rodents (As, Apr, Aal, A1-4, Intermediate, B) and in monkeys and humans (A_dark_, A_pale_, B1-4 (only one generation of B in human)), our comparative analysis of spermatogonia to transitioning preleptotene spermatocytes suggest that spermatogonial stem cells traverse through molecularly comparable states across species. Furthermore, we find that the proportion of cells contributed by each consensus state varies by species, reflecting inherent differences in the size of the stem/progenitor pool and the number of transit-amplifying divisions in the differentiating spermatogonia compartment across species. In these six states (SPG1-SPG6), we identify two molecularly distinct undifferentiated spermatogonial cell states (SPG1: TSPAN33, PIWIL4, CDK17, MORC1, ID4, ZBTB16, and SPG2: L1TD1, ID4, GFRA1, and ZBTB16). Immunohistochemistry analysis of these markers shows that SPG1 and SPG2 do not have a direct, one-to-one correspondence to the two histologically defined primate spermatogonial stem cell populations in the human testis known as A_dark_ or A_pale_. Rather, together SPG1 and 2 represent a heterogeneous pool of undifferentiated spermatogonia. Furthermore, we used human-to-nude mouse xenotransplantation to demonstrate that stem cell activity is enriched in the TSPAN33-positive fraction of SPG1, but was not restricted to that fraction as roughly half of total stem cell activity was recovered in the TSPAN33-negative fraction. Future studies using additional markers are needed to resolve the cellular heterogeneity, stem cell potential, and the phenotype(s) of the TSPAN33-negative stem cells. Based on the current data, it is not possible to know whether stem cell activity in the TSPAN33-negative fraction is from TSPAN33-negative cells in SPG1 or might also include cells in SPG2. Nonetheless, these data clearly indicate that the transplantable hSSCs are heterogeneous, at least with respect to the TSPAN33 cell surface marker.

Although SPG1 cells in our dataset mainly come from macaque and human samples, they do contain a small number of PIWIL4 (a.k.a. Miwi2) expressing cells from mouse. They account for <2% of mouse spermatogonia, too small a fraction to be detected in our earlier work. Curiously, Miwi2 knockout mice exhibit progressive loss of germ cells^90,91^. With a *Miwi2^tdtomato^* reporter, MIWI2 expression is detectable in a small number of A_single_ but is more prevalent in A_paired_ and A_aligned_ undifferentiated spermatogonia. These Miwi2-expressing cells possess SSC activity as demonstrated by transplantation^92^, and are required for efficient regeneration of spermatogenesis after injury in the adult mouse testis^93^. Together, these findings suggest that a rare population of early-stage undifferentiated SPG cells may exist in mouse that are analogous to SPG1 cells in primates. In mice, these cells may represent a rare population of As, Apr, Aal undifferentiated spermatogonia that were missed in our previous analysis due to its smaller number of cells.

The consensus SPG3-5 contains cells that are transitioning from undifferentiated (A_al_ mouse; A_d/p_ monkey and human) to differentiating (A1-4, Aintermediate and B spermatogonia mouse; B1-4 macaque; B human). We link these molecular states with morphometric states described across species and identify two molecularly distinct Type B molecular states. Further, SPG6 includes type spermatogonial cells transitioning to preleptotene spermatocytes based on the expression of conserved meiotic genes, such as PRDM9, ZCWPW1, and REC8. In addition to the known, classic meiotic genes, our dataset uncovers many primate-specific genes (i.e., lacking mouse orthologs) that are active in human and macaque SPG6, including the simian-specific Variably Charged (VC) gene family (VCX, VCX2, VCX3A, VCX3B)^94^, which is an X chromosome multi-copy gene family theorized to play a role in meiotic drive^94,95^. Recent studies have shown that VCX may be regulated by PRDM9^96^, further underscoring its potential role in meiosis. Although the function of this gene family is not yet directly tested, CNVs involving VCX have been associated with nonobstructive azospermia^97^. In addition to the VC-family, many primate-specific zinc-finger containing proteins peak in expression at the SPG6 stage. Unlike other transcriptional regulators, zinc-finger proteins have undergone bursts of duplication in vertebrates, including the primate lineage^98,99^, and are rapidly evolving (dN/dS >1)^100^, indicating they may have evolved specific functions in primate spermatogenesis. Specifically, KRAB-ZFPs are rapidly expanding and evolving ZF family^101^, and several lines of experimental evidence demonstrate that endogenous retro elements are the primary target sequences, suggesting this diversity may stem from dynamic competition between TEs and KRAB-ZFPs (reviewed in^102,103^). Based on the timing of expression of the primate-specific KRAB-ZF genes shown here, these genes may be activated in order to protect against the sequelae of active or inactive EREs, which are often also species-specific.

Taken together, our analysis suggests that many mammalian species traverse through comparable molecular states, which share some conserved features, while diverging on other features, reflecting evolving molecular pathways that execute the germ cell differentiation process.

Although the post-SPG portion of the spermatogenesis program (meiosis and spermiogenesis) is largely continuous, our multi-species comparisons of germ cell trajectories suggest that different species do not always follow the same progression through transcriptional intermediates during germ cell differentiation. Rather they exhibit heterochrony in the alignment of cellular states, both in the beginning (meiosis) or end (spermiogenesis) stages and in the pace of progression in the middle of the germ cell differentiation process. This apparent meandering in the comparison of germ cell trajectories is more pronounce with increasing evolutionary distance (human vs. mouse). To directly compare molecular states across species, we constructed a universal spermatogenesis “pseudotime” as the consensus of the three species and aligned molecularly analogous populations across species. This universal pseudotemporal map represents the first common molecular atlas of mammalian germ cell differentiation and provides state-specific markers in analogous populations in each species. With cells from all three species now mapped on the same scale, we were able to directly compare molecular states to reveal similarities and differences in the transcriptome and temporal shifts in gene expression. Interestingly, many of the genes experiencing shifted phase across species include transcription factors and RNA binding proteins.

Finally, we also discovered interspecies differences in the somatic cells that provide surrounding niche for the developing germ cells. The new catalog of somatic cells are accompanied by their molecular markers, expected functions, and likely communications among distinct cell types. A direct comparison of the somatic cell populations across species shows that the transcriptome of endothelial cells and immune cells are highly conserved across species, whereas other interstitial cells show greater divergence. For example, mouse myoid cells do not have the highest correlation to the human myoid cells, but instead bear a high transcriptional resemblance to human m-pericytes. To gain a preliminary view of the soma-germ cell communication and its potentially altered function across species, we examined known ligand-receptor pairs with predicted signaling from somatic cell populations to germ cells. We find that although certain pathways are likely employed to regulate similar stages of spermatogenesis, many other ligands/receptor pairs have either changed the source (cells of secretion) or the target (cells bearing the receptor), or both, generating many interesting hypotheses regarding the evolution of germ cell – soma communication in mammals. One example is the FGF signaling pathway which provides regulatory cues for spermatogonial dynamics^104,105^. FGF signaling depends on proteoglycans, such as syndecans, to bind and transfer FGFs to receptor tyrosine kinases (FGFRs) on target cells. In mouse spermatogonia, SDC4 promotes the consumption of FGFs^105^. We find that this interaction appears to be conserved in human and macaque, and there is an additional syndecan (SDC2) expressed in primates which can bind FGFs secreted by various somatic cells. Therefore, germline-soma communications have changed over the course of evolution through both the receptor and the ligand.

Altogether, our analyses provide the first reference system for integrated interspecies comparisons of spermatogenesis, highlighting multiple ways by which this process has evolved while maintaining the capacity to reach a common goal – male gamete production. Furthermore, we identify molecularly equivalent states between human, macaque, and mouse, so that despite the many apparent differences, mechanistic parallels can be drawn between species to enable future studies that make joint use of these complementary model systems. This knowledge enhances our ability to interpret contemporary discoveries in the context of the classic descriptions of cell morphology from the past. It is expected to reduce barriers to studying human infertility, and accelerate the progress towards developing novel therapies, such as *in vitro* gametogenesis.

## Acknowledgements

We thank members of the Hammoud, Li, Orwig, and Yamashita Labs for scientific discussions and manuscript comments. This research was supported by National Institute of Health (NIH) grants 1R21HD090371-01A1 (S.S.H. and J.Z.L.), 1DP2HD091949-01 (S.S.H.), P01 HD075795, R01 HD076412 (K.E.O), R01 HD092084 (K.E.O., S.S.H), F30HD097961 (A.N.S), training grants 5T32HD079342 (A.N.S.), 5T32GM007863 (A.N.S.), T32HD0871494 (S.K.M.), and Michigan Institute for Data Science (MIDAS) grant for Health Sciences Challenge Award (J.Z.L. and S.S.H.), Open Philanthropy Grant 2019-199327 (5384).

## Author contributions

S.S.H. and A.N.S. overall project design. J.Z.L. oversaw the computational analysis. K.E.O oversaw functional experiments. A.N.S., S.K.M., C.D.G., and M.S. performed experiments. X.Z., Q.M., and A.N.S. analyzed data. A.N.S and S.S.H. wrote the manuscript with input from J.Z.L. and K.E.O. Comments from all authors were provided.

