## Supplemental Figures 1-7 for "Single-cell RNA sequencing of human, macaque, and mouse testes uncovers conserved and divergent features of mammalian spermatogenesis"

### Figure S1

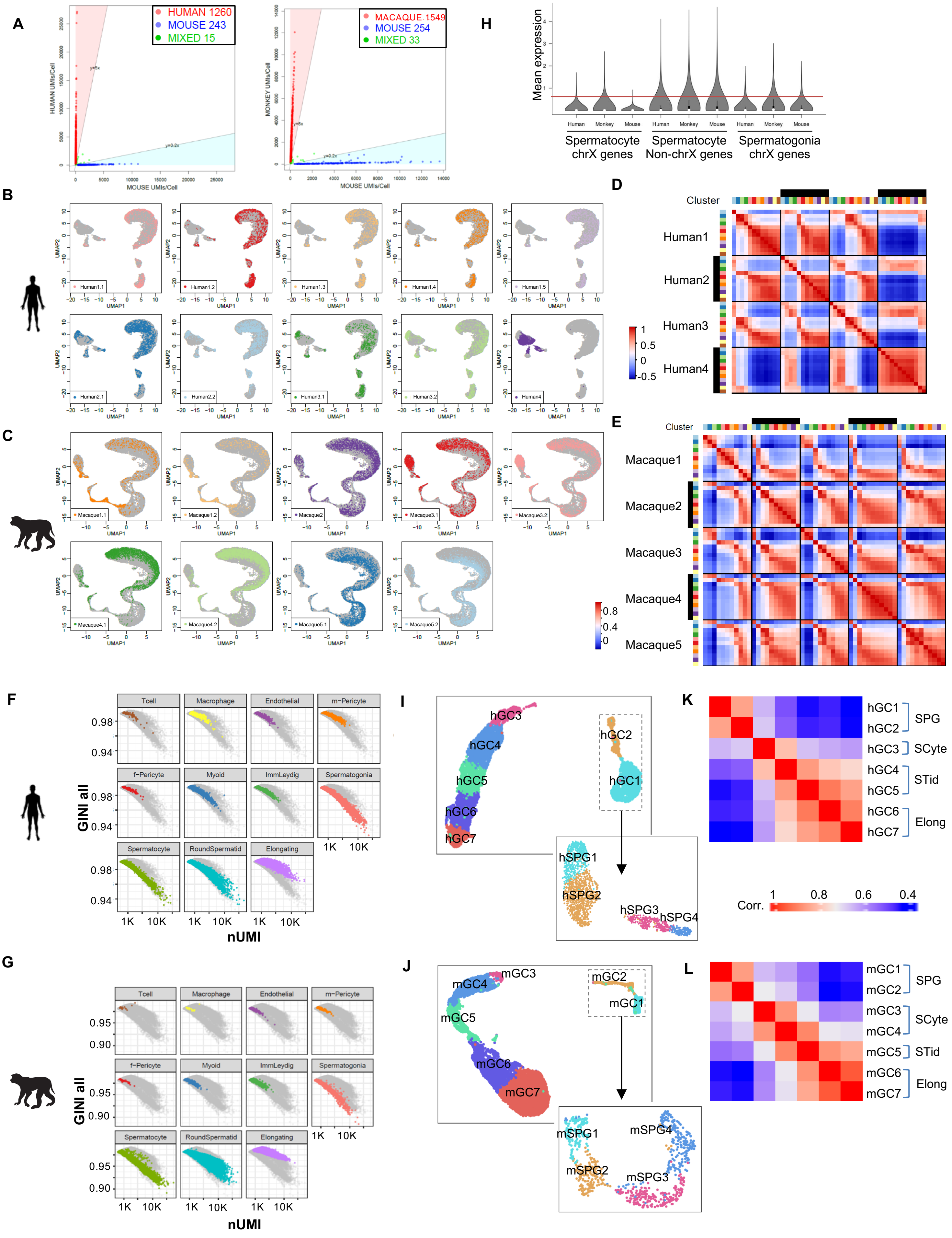
A
HUMAN 1260
MOUSE 243
MIXED 15
HUMAN UMIs/Cell
MOUSE UMIs/Cell
MACAQUE 1549
MOUSE 254
MIXED 33
MONKEY UMIs/Cell
MOUSE UMIs/Cell
H
Mean expression
Human
Monkey
Mouse
Spermatocyte chrX genes
Spermatocyte Non-chrX genes
Spermatogonia chrX genes
B
UMAP1
UMAP2
Human1.1
Human1.2
Human1.3
Human1.4
Human1.5
Human2.1
Human2.2
Human3.1
Human3.2
Human4
D
Cluster
Human1
Human2
Human3
Human4
C
Macaque1.1
Macaque1.2
Macaque2
Macaque3.1
Macaque3.2
Macaque4.1
Macaque4.2
Macaque5.1
Macaque5.2
E
Cluster
Macaque1
Macaque2
Macaque3
Macaque4
Macaque5
F
Tcell
Macrophage
Endothelial
m−Pericyte
f−Pericyte
Myoid
ImmLeydig
Spermatogonia
Spermatocyte
RoundSpermatid
Elongating
GINI all
nUMI
I
hGC1
hGC2
hGC3
hGC4
hGC5
hGC6
hGC7
hSPG1
hSPG2
hSPG3
hSPG4
K
SPG
SCyte
STid
Elong
Corr.
G
GINI all
nUMI
J
mGC1
mGC2
mGC3
mGC4
mGC5
mGC6
mGC7
mSPG1
mSPG2
mSPG3
mSPG4
L
SPG
SCyte
STid
Elong

### Figure S2

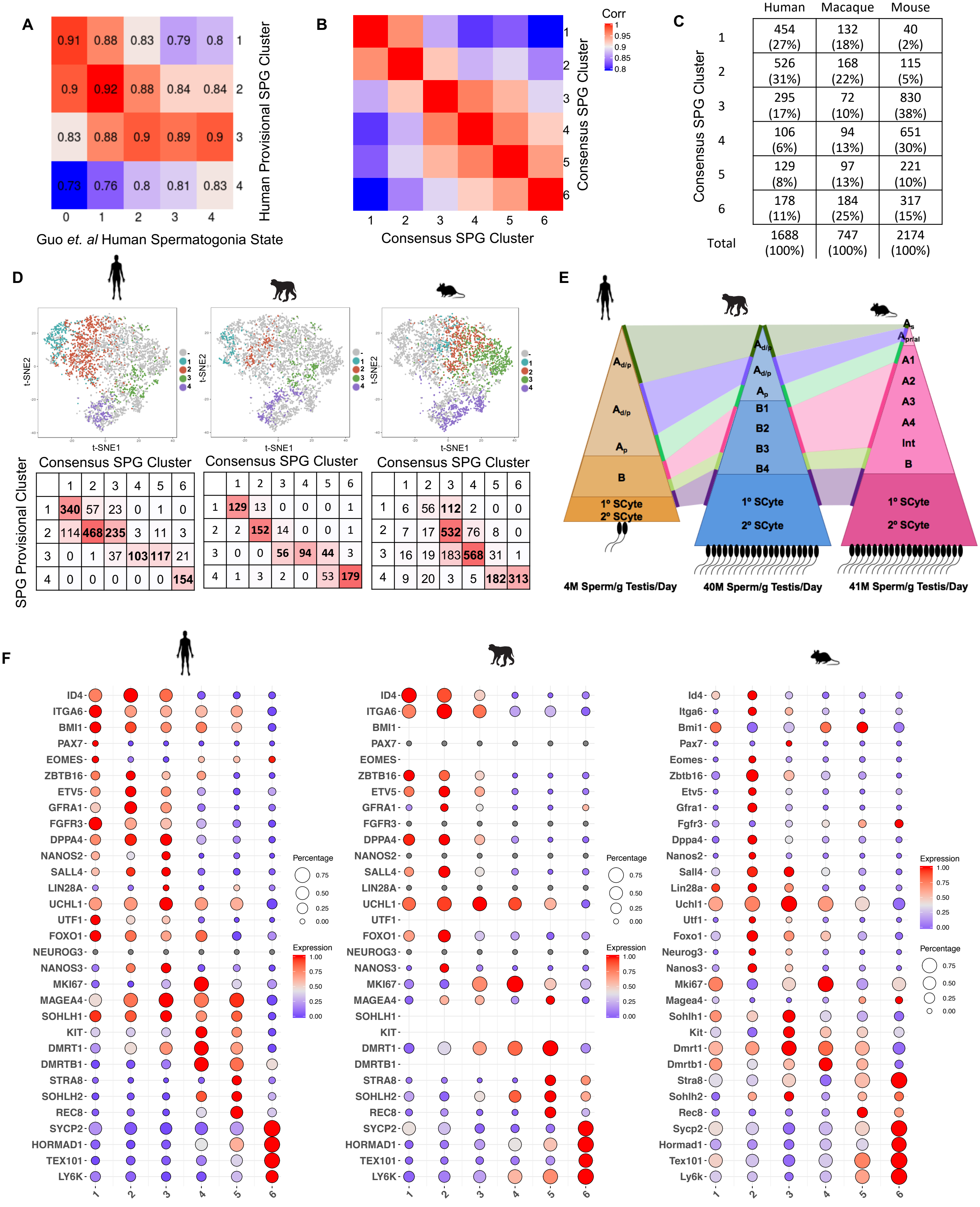
A
0.91 0.88 0.83 0.79 0.8 1
0.9 0.92 0.88 0.84 0.84 2
0.83 0.88 0.9 0.89 0.9 3
0.73 0.76 0.8 0.81 0.83 4
0 1 2 3 4
Guo et. al Human Spermatogonia State
Human Provisional SPG Cluster
B
Corr
1
0.95
0.9
0.85
0.8
1 2 3 4 5 6
Consensus SPG Cluster
Consensus SPG Cluster
C
Human Macaque Mouse
Consensus SPG Cluster
1 454 (27%) 132 (18%) 40 (2%)
2 526 (31%) 168 (22%) 115 (5%)
3 295 (17%) 72 (10%) 830 (38%)
4 106 (6%) 94 (13%) 651 (30%)
5 129 (8%) 97 (13%) 221 (10%)
6 178 (11%) 184 (25%) 317 (15%)
Total 1688 (100%) 747 (100%) 2174 (100%)
D
t-SNE1
t-SNE2
SPG Provisional Cluster
Consensus SPG Cluster
1 2 3 4 5 6
1 340 57 23 0 1 0
2 114 468 235 3 11 3
3 0 1 37 103 117 21
4 0 0 0 0 0 154
Consensus SPG Cluster
1 2 3 4 5 6
1 129 13 0 0 0 1
2 2 152 14 0 0 1
3 0 0 56 94 44 3
4 1 3 2 0 53 179
Consensus SPG Cluster
1 2 3 4 5 6
1 6 56 112 2 0 0
2 7 17 532 76 8 0
3 16 19 183 568 31 1
4 9 20 3 5 182 313
E
Ad/p
Ad/p
Ap
B
1° SCyte
2° SCyte
Ad/p
Ad/p
Ap
B1
B2
B3
B4
1° SCyte
2° SCyte
As
Apr/al
A1
A2
A3
A4
Int
B
1° SCyte
2° SCyte
4M Sperm/g Testis/Day
40M Sperm/g Testis/Day
41M Sperm/g Testis/Day
F
ID4 ITGA6 BMI1 PAX7 EOMES ZBTB16 ETV5 GFRA1 FGFR3 DPPA4 NANOS2 SALL4 LIN28A UCHL1 UTF1 FOXO1 NEUROG3 NANOS3 MKI67 MAGEA4 SOHLH1 KIT DMRT1 DMRTB1 STRA8 SOHLH2 REC8 SYCP2 HORMAD1 TEX101 LY6K
Id4 Itga6 Bmi1 Pax7 Eomes Zbtb16 Etv5 Gfra1 Fgfr3 Dppa4 Nanos2 Sall4 Lin28a Uchl1 Utf1 Foxo1 Neurog3 Nanos3 Mki67 Magea4 Sohlh1 Kit Dmrt1 Dmrtb1 Stra8 Sohlh2 Rec8 Sycp2 Hormad1 Tex101 Ly6k
1 2 3 4 5 6
Percentage
0.75
0.50
0.25
0.00
Expression
1.00
0.75
0.50
0.25
0.00

### Figure S3

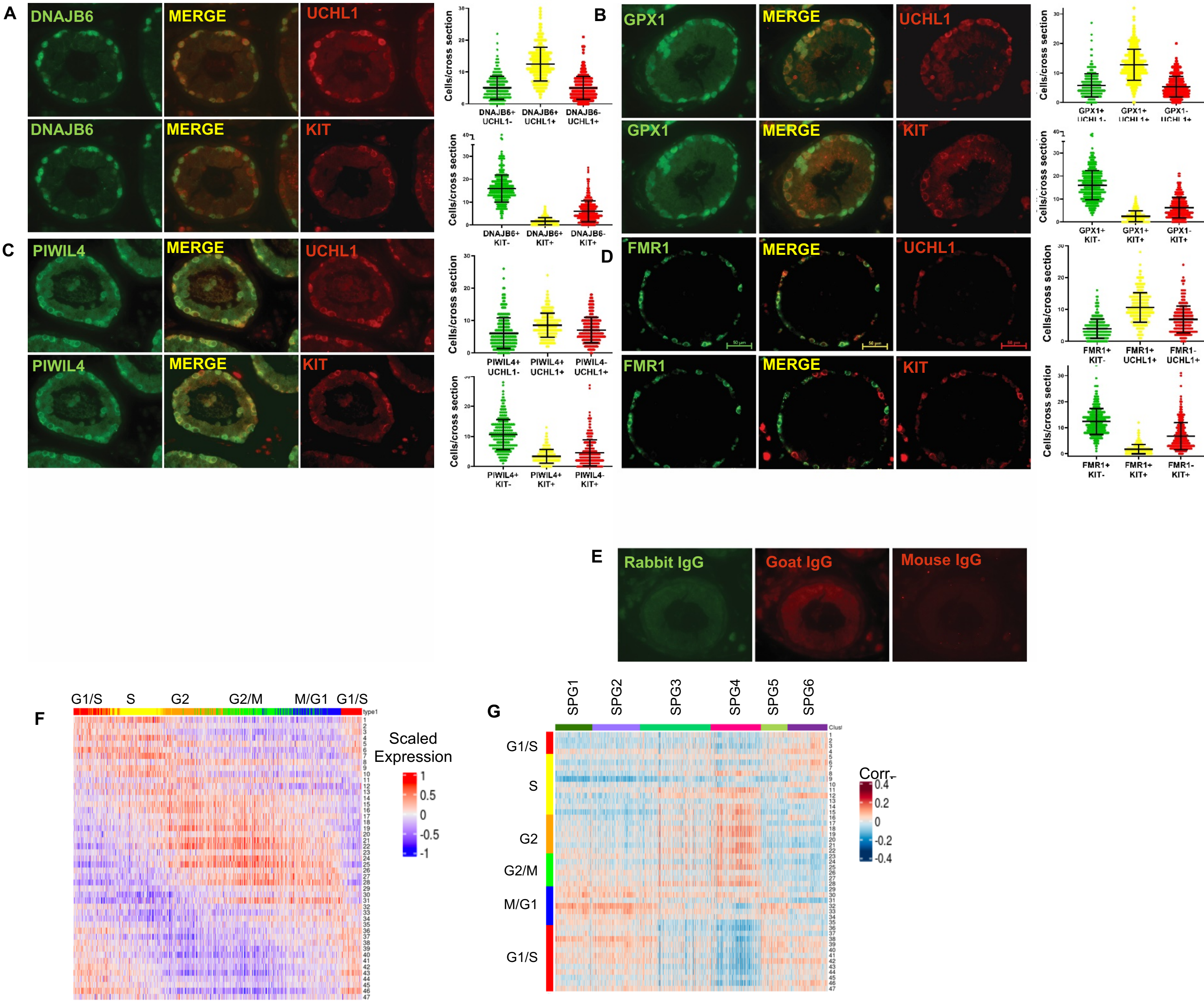


H

| | SPG1 | SPG2 | SPG3 | SPG4 | SPG5 | SPG6 |
|---|---|---|---|---|---|---|
| G2 | 31 (5%) | 12 (1.4%) | 114 (9.5%) | **284 (33.3%)** | 16 (3.6%) | 68 (10%) |
| G2/M | 66 (10.5%) | 49 (6.1%) | 208 (17.4%) | 158 (18.6%) | 26 (5.8%) | 11 (1.6%) |
| M/G1 | **365 (58.3%)** | **477 (59%)** | **339 (28.3%)** | 65 (7.6%) | **144 (32.2%)** | 58 (8.5%) |
| G1/S | **142 (22.7%)** | **247 (30.5%)** | **332 (27.7%)** | 99 (11.6%) | **213 (47.7%)** | **258 (40%)** |
| S | 22 (3.5%) | 24 (3%) | 204 (17%) | **245 (28.7%)** | 48 (10.7%) | **284 (41.8%)** |

### Figure S4

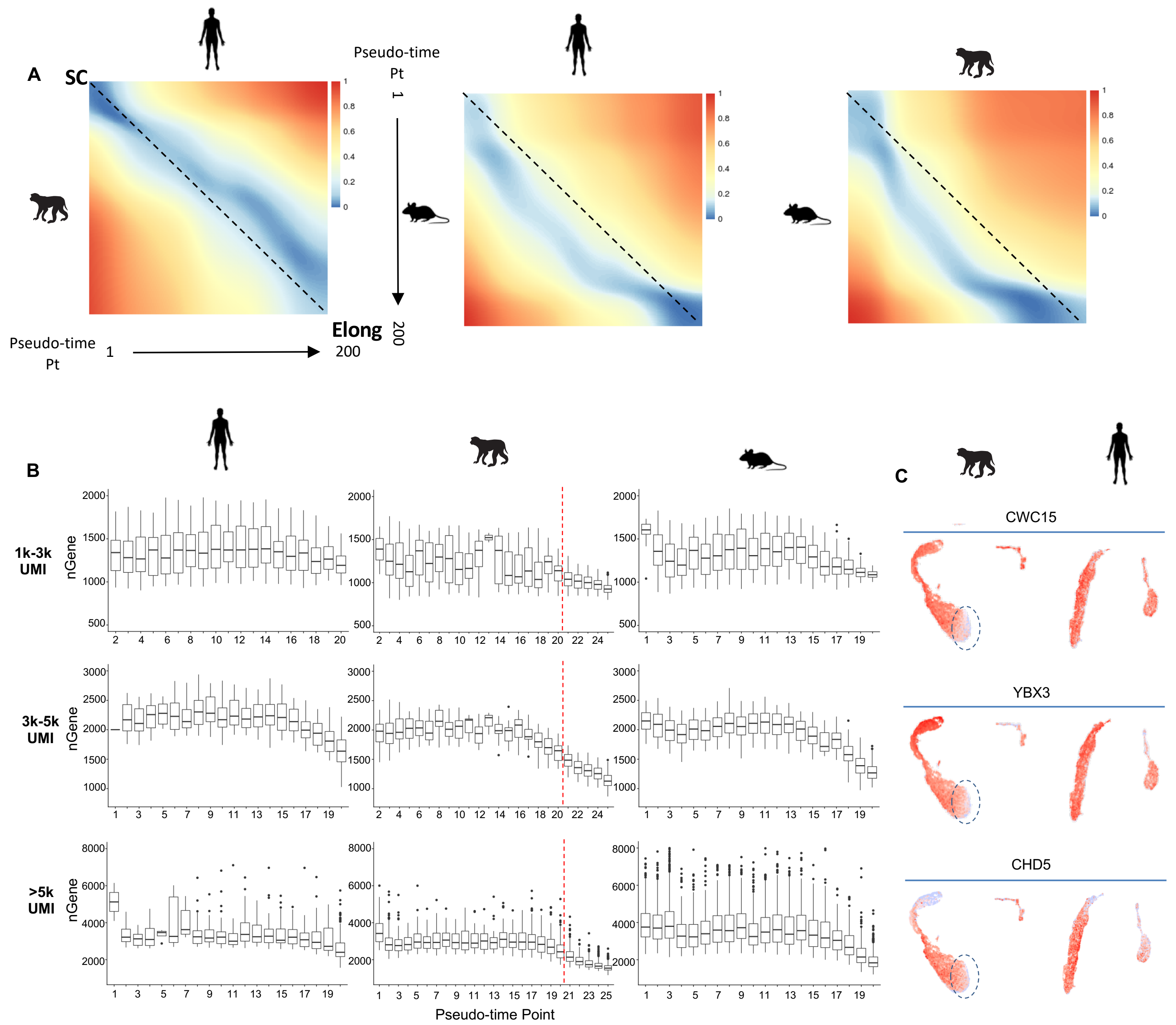

A
SC
Elong
Pseudo-time Pt
1
200
B
1k-3k UMI
3k-5k UMI
>5k UMI
nGene
Pseudo-time Point
C
CWC15
YBX3
CHD5

#### Figure S5

**A**

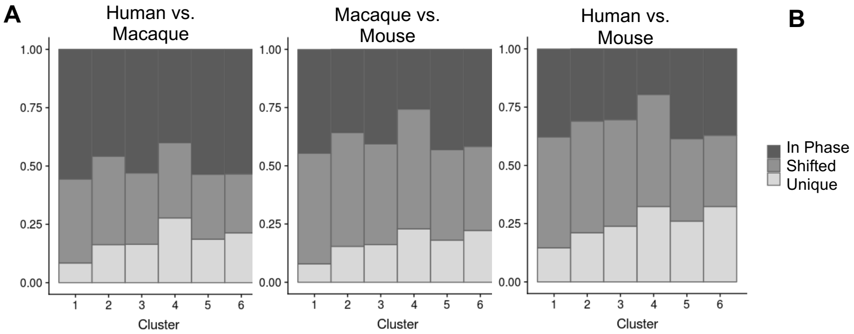


**B**

| Ortholog Type | | | Human | Macaque | Mouse |
|---|---|---|---|---|---|
| | α1 | Human specific | 324 | | |
| | α2 | Macaque specific | | 93 | |
| | α3 | Mouse specific | | | 2187 |
| | β1 | Human-Macaque specific | 875 | 862 | |
| | β2 | Mouse-Macaque specific | | 165 | 285 |
| | β3 | Human-Mouse specific | 1172 | | 1912 |
| | γ | x:y:z Multi-orthologs | 1131 | 2879 | 1744 |
| | δ | 1:1:1 orthologs | 12400 | 12078 | 13509 |
| | | All coding genes | 15902 | 16077 | 19637 |

**C**

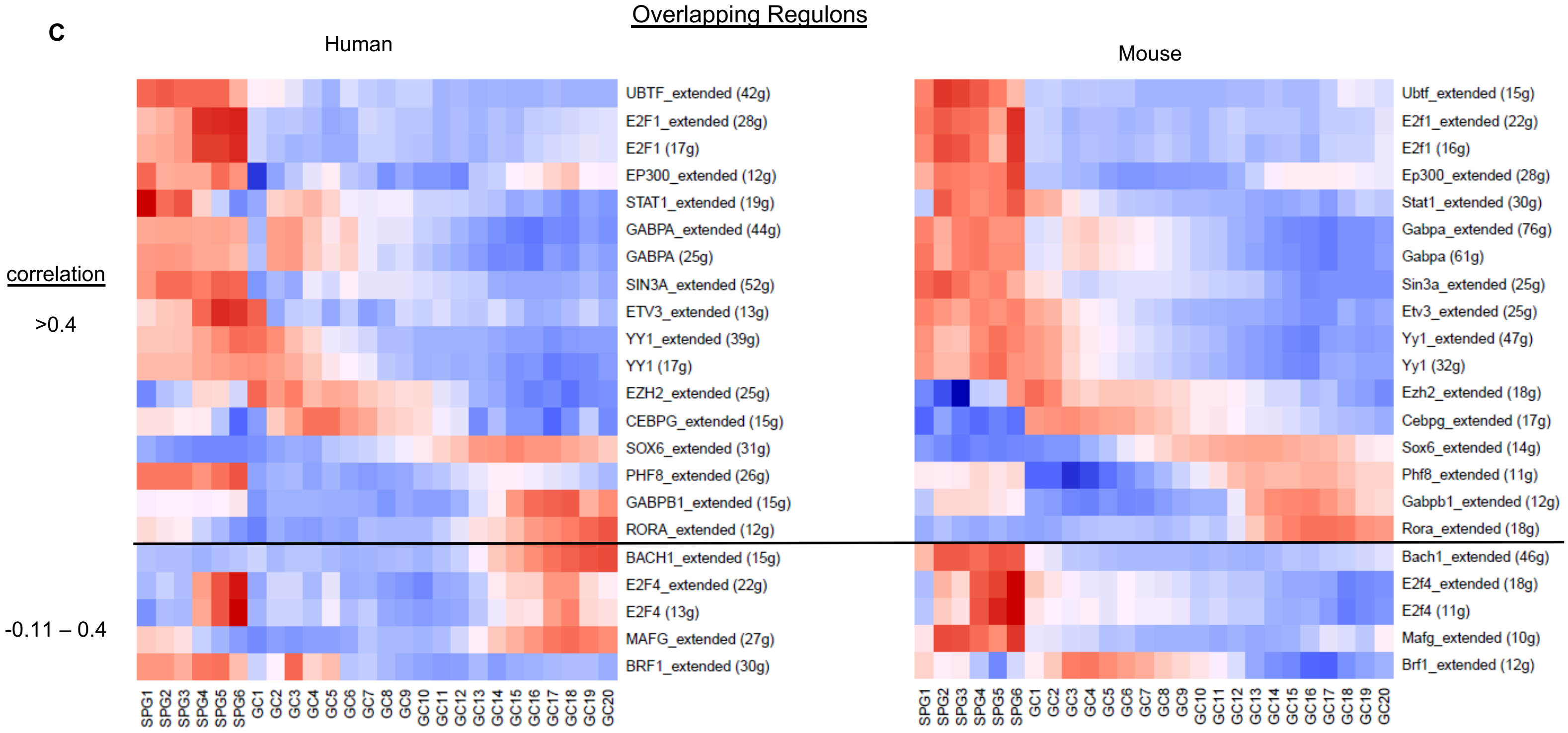


**D**

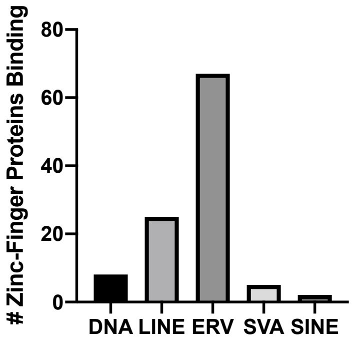


**E**

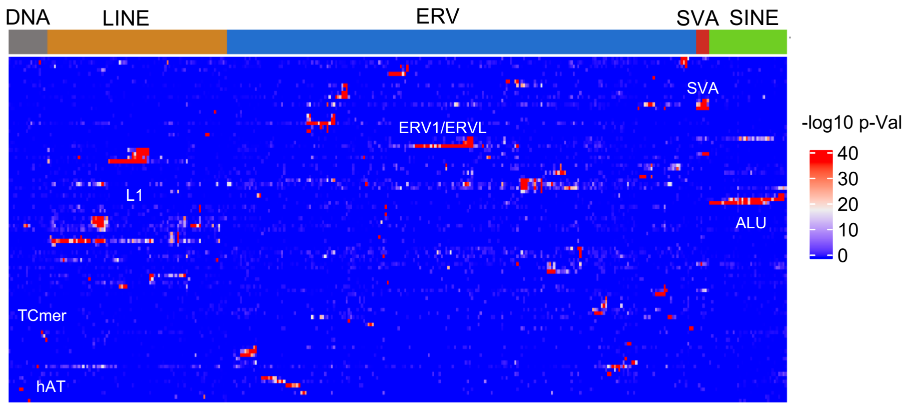

#### Figure S6

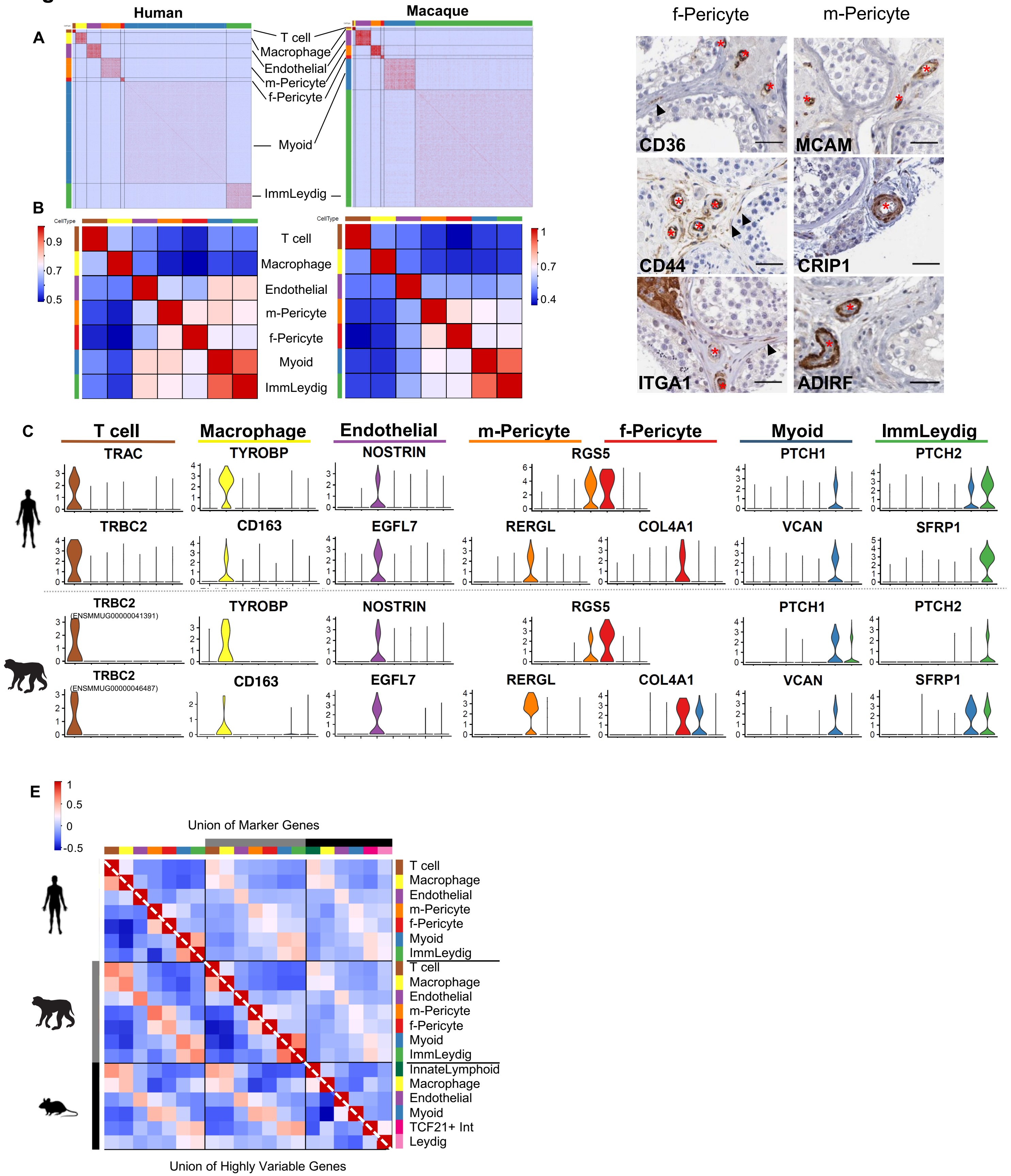

A
B
C
D
E
Human
Macaque
T cell
Macrophage
Endothelial
m-Pericyte
f-Pericyte
Myoid
ImmLeydig
CellType
0.9
0.7
0.5
1
0.7
0.4
TRAC
TYROBP
NOSTRIN
RGS5
PTCH1
PTCH2
TRBC2
CD163
EGFL7
RERGL
COL4A1
VCAN
SFRP1
TRBC2 (ENSMMUG00000041391)
TRBC2 (ENSMMUG00000046487)
f-Pericyte
m-Pericyte
CD36
MCAM
CD44
CRIP1
ITGA1
ADIRF
Union of Marker Genes
Union of Highly Variable Genes
0.5
0
-0.5
InnateLymphoid
TCF21+ Int
Leydig

### Figure S7

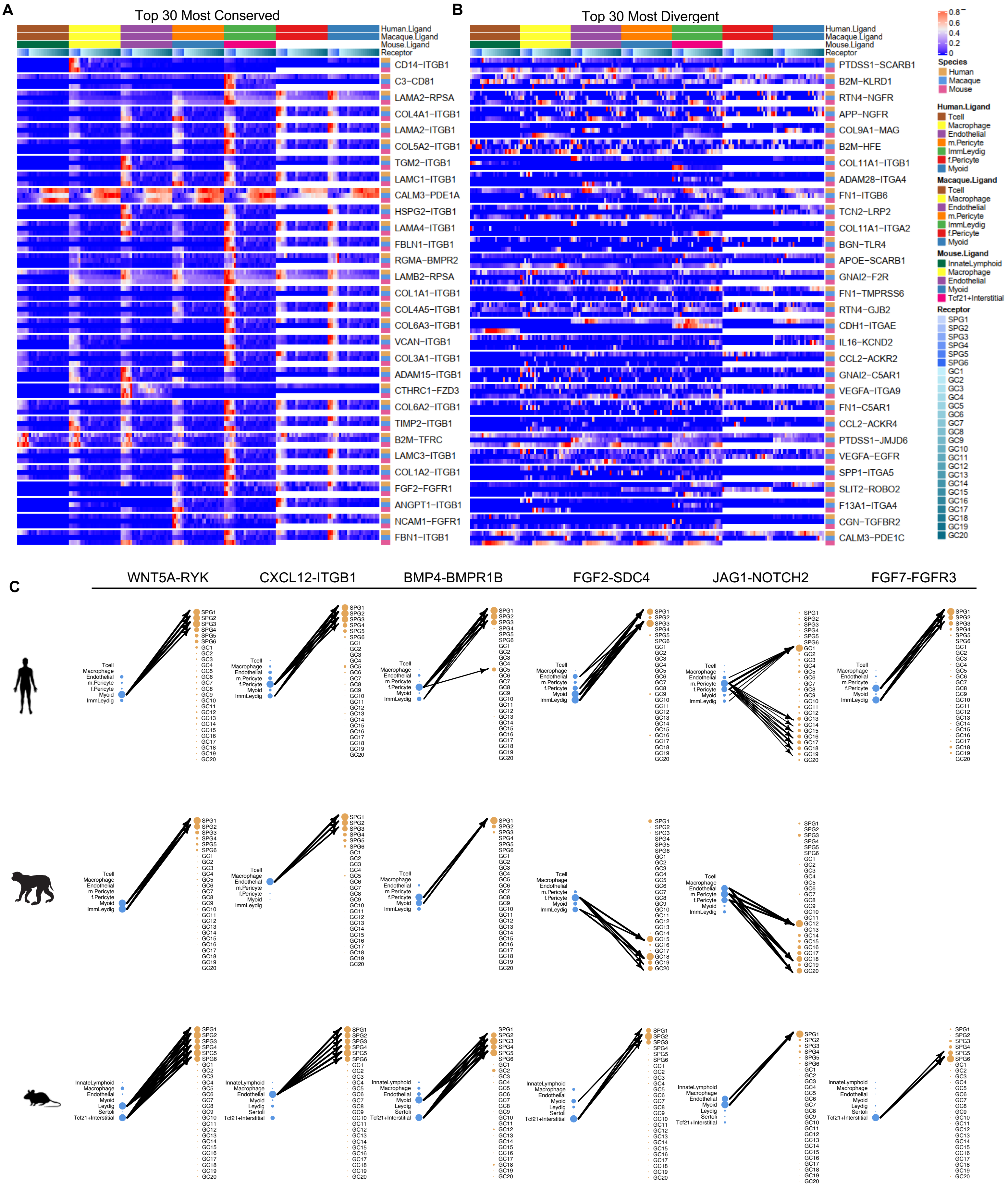
A
Top 30 Most Conserved
Human.Ligand
Macaque.Ligand
Mouse.Ligand
Receptor
CD14−ITGB1
C3−CD81
LAMA2−RPSA
COL4A1−ITGB1
LAMA2−ITGB1
COL5A2−ITGB1
TGM2−ITGB1
LAMC1−ITGB1
CALM3−PDE1A
HSPG2−ITGB1
LAMA4−ITGB1
FBLN1−ITGB1
RGMA−BMPR2
LAMB2−RPSA
COL1A1−ITGB1
COL4A5−ITGB1
COL6A3−ITGB1
VCAN−ITGB1
COL3A1−ITGB1
ADAM15−ITGB1
CTHRC1−FZD3
COL6A2−ITGB1
TIMP2−ITGB1
B2M−TFRC
LAMC3−ITGB1
COL1A2−ITGB1
FGF2−FGFR1
ANGPT1−ITGB1
NCAM1−FGFR1
FBN1−ITGB1
B
Top 30 Most Divergent
Human.Ligand
Macaque.Ligand
Mouse.Ligand
Receptor
PTDSS1−SCARB1
B2M−KLRD1
RTN4−NGFR
APP−NGFR
COL9A1−MAG
B2M−HFE
COL11A1−ITGB1
ADAM28−ITGA4
FN1−ITGB6
TCN2−LRP2
COL11A1−ITGA2
BGN−TLR4
APOE−SCARB1
GNAI2−F2R
FN1−TMPRSS6
RTN4−GJB2
CDH1−ITGAE
IL16−KCND2
CCL2−ACKR2
GNAI2−C5AR1
VEGFA−ITGA9
FN1−C5AR1
CCL2−ACKR4
PTDSS1−JMJD6
VEGFA−EGFR
SPP1−ITGA5
SLIT2−ROBO2
F13A1−ITGA4
CGN−TGFBR2
CALM3−PDE1C
0.8
0.6
0.4
0.2
0
Species
Human
Macaque
Mouse
Human.Ligand
Tcell
Macrophage
Endothelial
m.Pericyte
ImmLeydig
f.Pericyte
Myoid
Macaque.Ligand
Tcell
Macrophage
Endothelial
m.Pericyte
ImmLeydig
f.Pericyte
Myoid
Mouse.Ligand
InnateLymphoid
Macrophage
Endothelial
Myoid
Tcf21+Interstitial
Receptor
SPG1
SPG2
SPG3
SPG4
SPG5
SPG6
GC1
GC2
GC3
GC4
GC5
GC6
GC7
GC8
GC9
GC10
GC11
GC12
GC13
GC14
GC15
GC16
GC17
GC18
GC19
GC20
C
WNT5A-RYK
CXCL12-ITGB1
BMP4-BMPR1B
FGF2-SDC4
JAG1-NOTCH2
FGF7-FGFR3
Tcell
Macrophage
Endothelial
m.Pericyte
f.Pericyte
Myoid
ImmLeydig
InnateLymphoid
Leydig
Sertoli
Tcf21+Interstitial
SPG1
SPG2
SPG3
SPG4
SPG5
SPG6
GC1
GC2
GC3
GC4
GC5
GC6
GC7
GC8
GC9
GC10
GC11
GC12
GC13
GC14
GC15
GC16
GC17
GC18
GC19
GC20
